# Towards Stewardship of Wild Species and Their Domesticated Counterparts: A Case Study in Northern Wild Rice (*Zizania palustris* L.)

**DOI:** 10.1101/2022.08.25.505308

**Authors:** Lillian McGilp, Matthew W. Haas, Mingqin Shao, Reneth Millas, Claudia Castell-Miller, Anthony J. Kern, Laura M. Shannon, Jennifer A. Kimball

**Author notes:** Department of Energy Joint Genome Institute, Lawrence Berkeley National Laboratory, Berkeley, CA 94720, USA.

## Abstract

Northern Wild Rice (NWR; *Zizania palustris* L.) is an aquatic, annual grass with significant ecological, cultural, and economic importance to the Great Lakes region of North America. In this study, we assembled and genotyped a diverse collection of 839 NWR individuals using genotyping-by-sequencing (GBS) and obtained 5,955 single-nucleotide polymorphisms (SNPs). Our collection consisted of samples from 12 wild NWR populations collected across Minnesota and Western Wisconsin, some of which were collected over two time points; a representative collection of cultivated NWR varieties and breeding populations; and a *Zizania aquatica* outgroup. Using these data, we characterized the genetic diversity, relatedness, and population structure of this broad collection of NWR genotypes. We found that wild populations of NWR clustered primarily by their geographical location, with some clustering patterns likely influenced by historical ecosystem management. Cultivated populations were genetically distinct from wild populations, suggesting limited gene flow between the semi-domesticated crop and its wild counterparts. The first genome-wide scans of putative selection events in cultivated NWR suggest that the crop is undergoing heavy selection pressure for traits conducive to irrigated paddy conditions. Overall, this study presents a large set of SNP markers for use in NWR genetic studies and provides new insights into the gene flow, history, and complexity of wild and cultivated populations of NWR.

## Introduction

Northern Wild Rice (NWR; *Zizania palustris* L.) is an annual, diploid (2*n*=2*x*=30), aquatic grass native to the Eastern Temperate and Northern Forest ecoregions of North America ^1,2^. Predominantly found in shallow, slow-moving waters surrounding the Great Lakes region of North America, NWR’s regional significance in these areas is vast and complex ^3–8^. To begin, NWR is a natural resource that provides food and substantial habitat for a wide range of wildlife ^9,10^ as well as important ecosystem services such as anchoring riparian soils and inhibiting algal blooms ^11^. For centuries, this nutritious grain has been hand-harvested from regional lakes and rivers by Dakota and Anishinaabe Peoples ^2,12^, and ‘psin’ (Dakota) or ‘manoomin’ (Ojibwe, Anishinaabe) remains an integral component of their cultures and lives today. The species has also become a high-value commodity crop that includes hand-harvested grain from regional waters, and cultivated grain from irrigated paddies, grown primarily in Minnesota (MN) and California (CA) ^13^. As a part of the *Oryzae* tribe in the *Poaceae* family, *Zizania* species are also considered crop wild relatives of *Oryza sativa* L. (white rice). Given the many roles described above, we contend that the conservation of NWR serves as an important intersection between our ecosystems, our cultures and food, and our economies.

Worldwide, plant species are experiencing declines and extinction events as a result of human activities altering natural ecosystems. For at least the last century, NWR has been experiencing such declines in its native habitats ^14,15^, and the species appears to be slowly migrating northward ^16^. Recent reports have stated that NWR is at high risk of loss, and the International Union for Conservation of Nature has added NWR to their red list of threatened species ^17^. Hydrological changes due to damming and channelization, recreational water activity, shoreline development, and water pollution from industrial activities have all been associated with the decline of NWR in its natural habitats ^6,15^. The species is particularly sensitive to high levels of sulfates in its water supply and acts as an important indicator species of water quality ^6,18^. In addition to these conservation challenges, NWR seed is intermediately recalcitrant or desiccation intolerant, which reduces seed longevity in storage to 1-2 years ^19^ and reduces the feasibility of preservation in *ex-situ* seed banks. Therefore, the genetic diversity of NWR is currently only preserved *in-situ* within its natural range (Porter, 2019).

Genetic diversity represents the extent of heritable variation within and among populations of a species, and its preservation is vital for the maintenance of long-term viability in the face of continual environmental change ^20,21^. A 2008 MN Department of Natural Resources (MN DNR) report on the health of NWR natural stands concluded that the species’ greatest threat was an overall state-wide decline in genetic diversity ^15^. Biologists and conservationists have widely recognized the value of characterizing genome-wide diversity within a species for use in conservation efforts ^22,23^. However, few such studies have been conducted for NWR, and molecular studies have heavily relied on obsolete marker systems ^5,24,25^, which are laborious to produce and often limited in number ^26^. The advent of low-cost high-throughput sequencing, such as genotyping-by-sequencing (GBS), which provides genome-wide coverage of co-dominant, single-nucleotide polymorphism (SNP) markers, has improved the genetic diversity characterization of extensive and complex germplasm collections ^27^. In 2019, a study with limited sample size demonstrated the potential for GBS to be applied to NWR ^28^.

The cultivation of NWR in irrigated man-made paddies, similar to white rice production, began in the 1950s to create an industry capable of supplying a consistent source of the grain to agricultural markets. As such, the production of cultivated NWR (cNWR) is a fairly new endeavor and only ~60 cycles of targeted selection separate cNWR from its wild counterparts. Breeders of cNWR have focused primarily on adapting the species to agronomic production in paddies, and on fixing seed-shattering resistance in the crop ^2^. However, concerns regarding the potential impact of gene flow between cNWR and natural stands of NWR have been raised given the species’ outcrossing nature ^12,29–31^. The Great Lakes region is the center of both origin and diversity of *Z. palustris*; therefore, it is important to understand the potential impact of domesticating and cultivating cNWR in those areas. As plant breeders, we have a responsibility to be good stewards of our natural and domesticated plants. Thus, an understanding of how cNWR fits into local landscapes, and particularly, to what extent gene flow is occurring among natural stands and cNWR is essential to overall germplasm preservation.

In this study, we generated a genome-wide SNP dataset via GBS for a NWR diversity collection to study the population structure and gene flow within and among wild and cultivated populations. We aimed to improve our understanding of the genetic variability within the species and provide new information regarding the selective pressures applied to cNWR germplasm.

## Materials and Methods

### Plant materials

The diversity collections consisted of wild populations gathered across northern MN and Wisconsin (WI) (referred to here as the Natural Stand collection); cultivars and germplasm from the UMN cNWR breeding program (referred to here as the Cultivated collection); and a population of *Zizania aquatica* L. from the Platte River in MN (referred to here as the outgroup). A Temporal panel was also generated to look at potential changes in diversity over time within two natural stand lake populations. In all, a total of 889 individuals were evaluated in this study, which consisted of 530 Natural Stand and 209 Cultivated samples collected in 2018, in addition to 100 samples collected in 2010 for the Temporal panel.

The Natural Stand collection was obtained from 10 wild populations across northern Minnesota (50 samples per lake/river) and 2 wild populations from western Wisconsin (10 from Mud Hen Lake; 20 from Phantom Lake) (Figure 1; Table S1). An additional 50 *Z. aquatica* samples, also collected in central Minnesota, were used as an outgroup in this study. Three major hydrologic unit code (HUC) subbasins were represented in this collection including seven populations from the Upper Mississippi River (UMR) watershed, three populations from the Red River of the North (RRN) watershed, and two populations from the St. Croix River (SCR) watershed (Figure 1; Table S1). A distance (km) matrix for all Natural Stand populations is available in Table S2.

**Figure 1.**
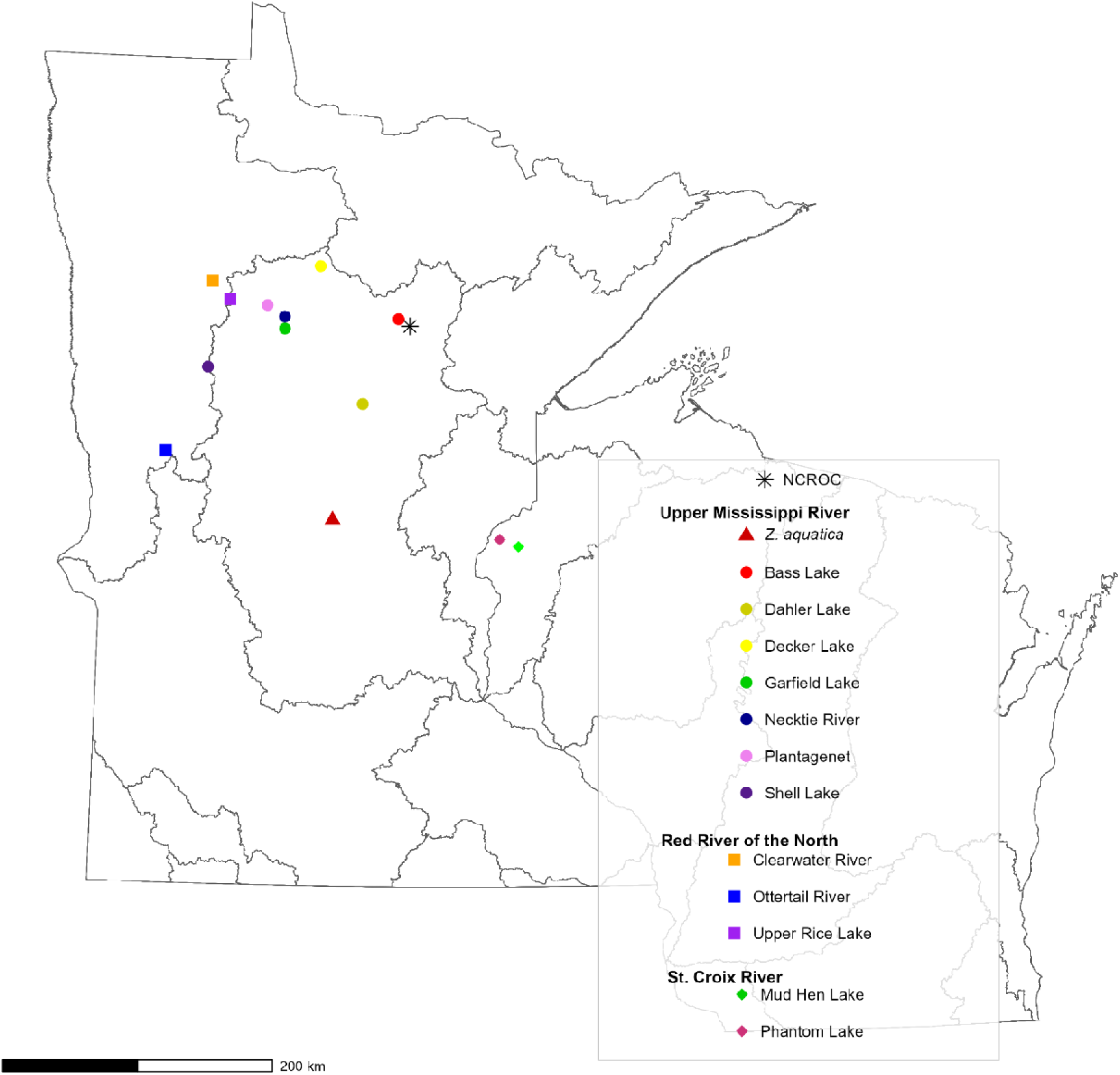
A Hydrological Unit Code-8 (HUC-8) watershed map of Minnesota and western Wisconsin indicating areas of sample collection of Northern Wild Rice (NWR; *Zizania palustri* L.) and *Zizania aquatica* L. GPS coordinates can be found in Table S1.

The Cultivated collection consisted of leaf samples from 209 open-pollinated individuals, representing 4 cultivars and 10 breeding populations from the UMN cNWR breeding program. Samples were collected at the UMN North Central Research and Outreach Center (NCROC) in Grand Rapids, MN in 2018 (Table S1; Figure 1; Figure S1).

Our Temporal collection consisted of 200 natural stand samples collected from Garfield and Shell Lakes in 2010 and in 2018, with 50 samples collected in each lake at each time point. GPS coordinates were used to ensure that the same population at the same location were collected at both time points (Table S1). We would like to note that samples from Garfield and Shell Lakes collected in 2018 were also a part of the Natural Stand collection (50 samples per lake).

### DNA extraction and sequencing

During collection, approximately 8 cm of leaf tissue was harvested on ice and then lyophilized using a TissueLyser II (Qiagen, Valencia, CA, USA). Genomic DNA was extracted using a Qiagen DNeasy Plant Mini Kit (Qiagen, Valencia, CA, USA) according to the manufacturer’s instructions. DNA concentration was measured with a NanoDrop spectrophotometer (Thermo Scientific, Wilmington, DE, USA), samples were submitted to the UMN Genomics Center for library preparation and sequencing. The digestion step was performed using two restriction enzymes, *Btg1* (5’-C/CRYGG-3’) and *TaqI* (5’-T/CGA-3’) (Shao et al., 2019). Afterward, unique barcodes were ligated to DNA fragments for sample identification and pooling. Single-end 150-bp sequencing to a depth of 2.5 million reads per sample was performed on an Illumina NovaSeq machine (Illumina, San Diego, CA, USA). Raw data were deposited in the National Center for Biotechnology Information Short Read Archive (NCBI SRA) under accession number PRJNA774842. BioSample accession numbers for individual samples are provided in Table S3.

### Read mapping

Quality control was initially performed on the FASTQ files using fastQC version 0.11.7 ^32^ to check for read quality and adapter contamination. After adapter trimming with Cutadapt version 1.18 ^33^, reads were mapped to the reference genome v1.0 of cNWR cultivar ‘Itasca-C12’ ^34^ of BWA-MEM version 0.7.13 ^35^. SNPs were called using the ‘mpileup’ function from BCFtools version 2.3 ^36^ and sorted with the sort function from samtools version 1.9 ^37^, resulting in 2,183 variant call format (VCF) files, one for each scaffold of the NWR genome ^34^. The 17 largest VCF files representing the 15 NWR chromosomes and 2 additional large scaffolds of the NWR reference genome were merged into a single VCF file with the ‘concat’ function from BCFtools. Filtering of the merged VCF file was carried out using VCFtools ^38^. All analyses were done with default parameters. A maximum missing rate of 20% across all samples and a minimum depth of 4 reads per variant site was used to obtain the final SNP set.

### Marker statistics

Summary statistics were calculated for each SNP including their distribution across chromosomes and scaffolds, their polymorphism information content (PIC) value, and their transition/transversion (TsTv) ratio. Polymorphism information content (PIC) was calculated using the *snpReady* R package ^39^. The number and type of transitions and transversions were calculated using VCFtools.

### Genetic Diversity Assessments

To assess the structure and distribution of genetic variation within our collections, we conducted a principal coordinate analysis (PCoA). To generate the full sample PCoA, the VCF file resulting from our SNP calling pipeline was imported into the R statistical environment ^40^ using the *vcfR* package ^41^. For the individual Natural Stand, Cultivated, and Temporal PCoAs, the original VCF file was first subsetted to only include the relevant samples using PLINK version 1.90b6.10 ^42^. For all PCoAs, the R package *vegan* ^43^ was used to calculate a dissimilarity matrix based on the Jaccard distance using the ‘vegdist’ function, as well as eigenvectors and eigenvalues using the ‘cmdscale’ function. PCoA plots were then generated using the ggplot2 package ^44^.

Neighbor-joining trees for the Natural Stand and Cultivated collections as well as the Temporal collection were created using the R packages *adegenet* ^45,46^, *ape* ^47^, *poppr* ^48,49^, and *vcfR*. The temporal cluster analysis was performed similarly to the methods described in Jacquemyn et al., 2006 ^50^. We estimated Prevosti’s genetic distance using the unweighted pair group method with arithmetic mean (UPGMA) algorithm option for the ‘aboot’ function (bootstrapped dendrograms) in *poppr* with the 1,000 bootstrap replicates, and a selected cutoff value of 50.

Population structure and admixture in our Natural Stand and Cultivated collections was assessed using Bayesian clustering implemented in STRUCTURE version 2.3.4 ^51^. Genotypic data for these individuals and the final bi-allelic SNP set were loaded into STRUCTURE and analyzed with the admixture model. The Markov Chain Monte Carlo was run from *K*=2 to *K*=14 with a burn-in length of 1,000 followed by 10,000 iterations. Lower *K* values were used to look for larger structural diversity patterns, while the higher *K* values were chosen to test if we were able to separate each population into their own STRUCTURE-assigned cluster (e.g., area of collection). Each *K* value was run 3 times, then compiled into a merged file using the ‘clumppExport’ function from the *pophelper* R package and subsequently plotted with the *pophelper package* ^52^. The ideal number of clusters was determined using the DeltaK statistic from the Structure Harvester web tool version 0.6.94 ^53^, which uses the Evanno et al. (2005) method for determining the number of clusters ^54^.

Analysis of Molecular Variance (AMOVA) was performed using the ‘poppr.amova’ function from the *poppr* R package based on the work of Excoffier et al. (1992) ^55^. Groups were defined based on their collection membership (e.g., their lake/river of origin or cultivar/breeding line identity), species (*Z. palustris* or *Z. aquatica*), and, more broadly, their germplasm type (e.g., Natural Stand or Cultivated). We performed AMOVAs using the “farthest neighbor” algorithm and the “quasieuclid” (default) method in the *ade4* package using default parameters ^56^. The significance was determined using the ‘randtest’ function with 999 repetitions.

Correlation between geographic and genetic distances in our Natural Stand collection, were assessed with a Mantel test using the ‘mantel.test’ function from the *ape* R package, based on genetic distances calculated with the ‘dist.genpop’ function from the *adegenet* R package and geographic distances calculated using the ‘distm’ function from the *geosphere* R package ^57^. The regression equation was found by fitting the genetic and geographic distances to a linear model.

### Gene flow

Pairwise estimations of genetic differentiation (*F_ST_*) ^58^ between different subgroups based on geographic origin and germplasm type (cultivated population, natural stand, species) were calculated using the ‘stamppFst’ function from the *StAMPP* R package ^59^. Dsuite version 0.4 ^60^ was used to calculate Patterson’s *D*-statistics, also known as the ABBA-BABA statistic, in order to test for introgressions between groups. Since this analysis requires four groups, we defined the groups based on their membership in STRUCTURE-assigned groups when *K*=4. This approach split the Natural Stands into two groups, while the Cultivated collection formed a third group. The fourth group (the outgroup) was *Z. aquatica*.

### Genome-wide Scans for Signatures of Selection

We used a series of tests to identify putative selection events in cultivated NWR. We calculated nucleotide diversity (π) ^61^, Tajima’s D ^62^, and F_ST_ ^58^ for each SNP using VCFtools. Natural Stand and Cultivated collections were analyzed separately. The results were plotted in the R statistical environment by averaging π values of ~10 Mb/bin. We also used the Cross-Population Composite Likelihood Ratio (XP-CLR) test ^63^ to test for large deviations between Natural Stand and Cultivated collections.

## Results

### Genotyping-by-sequencing

Using GBS technology, 839 *Z. palustris* samples representing the Natural Stand, Cultivated, and Temporal collections along with 50 samples of *Z. aquatic*a, which served as an outgroup, were sequenced to a depth of 2.5 million reads per sample. A total of 1,833,504,458 reads were generated with an average of 2,185,345 reads per sample. From these data, a total of 5,955 SNP markers were identified that met our filtering criteria, 3,005 (50.1%), of which, reside in genes. Basic marker statistics are shown in Table 1. The SNP density ranged from 1.15 - 6.37 SNPs/Mb per chromosome with a genome-wide average of 4.34 SNPs/Mb using 1Mb bins. In genic regions, the SNP density ranged from 10-20 SNPs/Mb per chromosome. The genome-wide TsTv ratio was 3.15 with a minimum of 1.50 (ZPchr0458) and a maximum of 4.18 (ZPchr0016) (Table S4). PIC ranged from 0.030 - 0.313, with an average of 0.217 (Table S5).

**Table 1.** Summary marker statistics for 5,955 Northern Wild Rice (NWR; *Zizania palustris* L.) bi-allelic single nucleotide polymorphism (SNP) markers generated via genotyping-by-sequencing (GBS).

| Chr/Scaffold | Size (Mbp) | SNPs (#) | % of all SNPs | SNPs in genic regions (#) | % of SNPs in genic regions/ Chr |
| --- | --- | --- | --- | --- | --- |
| ZPchr0001 | 95.4 | 483 | 8.11 | 74 | 15.32 |
| ZPchr0002 | 103.4 | 435 | 7.30 | 56 | 12.87 |
| ZPchr0003 | 58.8 | 302 | 5.07 | 37 | 12.25 |
| ZPchr0004 | 98.7 | 491 | 8.25 | 24 | 4.89 |
| ZPchr0005 | 66.6 | 350 | 5.88 | 41 | 11.71 |
| ZPchr0006 | 118.0 | 598 | 10.04 | 126 | 21.07 |
| ZPchr0007 | 42.6 | 181 | 3.04 | 66 | 36.46 |
| ZPchr0008 | 75.7 | 367 | 6.16 | 38 | 10.35 |
| ZPchr0009 | 95.1 | 481 | 8.08 | 25 | 5.20 |
| ZPchr0010 | 111.4 | 568 | 9.54 | 75 | 13.20 |
| ZPchr0011 | 63.2 | 285 | 4.76 | 41 | 14.39 |
| ZPchr0012 | 105.9 | 410 | 6.88 | 86 | 20.98 |
| ZPchr0013 | 111.3 | 583 | 9.79 | 66 | 11.32 |
| ZPchr0014 | 24.0 | 91 | 1.53 | 28 | 30.77 |
| ZPchr0015 | 39.1 | 237 | 3.98 | 12 | 5.06 |
| ZPchr0016* | 13.8 | 88 | 1.48 | 6 | 6.82 |
| ZPchr0458* | 4.3 | 5 | 0.08 | 3 | 60.00 |

\* Additional unaligned scaffolds from the NWR reference assembly v1.0 (Haas et al., 20201)

### Genetic Diversity of NWR Collections

To visualize the variation within the diversity collection, we first performed PCoA on both the Natural Stand and Cultivated collections together (Figure 2a). The first three principal coordinates explained 12.5%, 7.7%, and 7.3% of the variance, respectively (27.5% total). Within the first coordinate, samples were primarily split into two clusters, including a cluster of all *Z. aquatica* samples and the majority of Natural Stand samples, and a second cluster including all Cultivated samples and Bass and Decker Lake populations from the UMR watershed (Figure 2a). Upper Rice Lake genotypes appeared to blend between these two main groups but trended more heavily toward the group containing the UMR watershed genotypes. Within the first two principal coordinates, a large range of continuous variation can be seen, with samples primarily grouped by their germplasm type and geographic origin. Overall, these two principal coordinates failed to fully separate individual populations, though most *Z. aquatica* samples did separate from those of *Z. palustris* (Figure 2a). The Cultivated collection largely formed its own group with only a small number of samples overlapping with those from Bass, Upper Rice, Decker, Dahler, and Phantom (WI) Lakes. The third principal coordinate, in conjunction with coordinate 1, isolated *Z. aquatica* from all *Z. palustris* samples, Natural Stand and Cultivated collections, while maintaining the separation of the two main groups defined by coordinates 1 and 2 (Figure S2).

**Figure 2.**
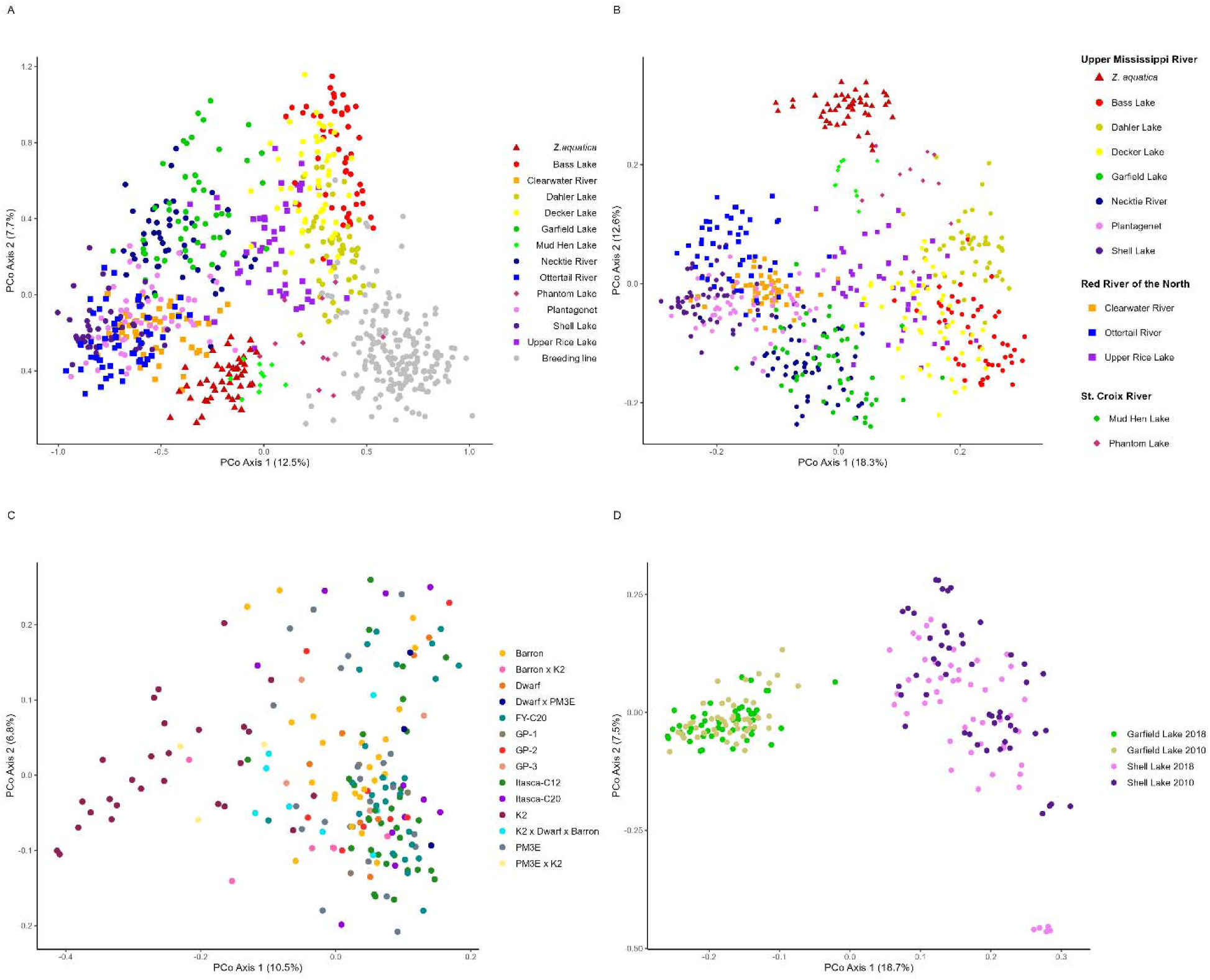
Principal coordinate analysis (PCoA) showing the differentiation of the 1^st^ and 2^nd^ principal coordinates of a.) the Natural Stand and Cultivated collections of Northern Wild Rice (NWR; *Zizania palustris* L.); b.) the Natural Stand collection; c.) the Cultivated collection; and d.) the Temporal collection.

When analyzed separately from the Cultivated collection, the first two coordinates of the Natural Stand collection explained 19% and 13% of the variance, respectively (Figure 2b). The two main clusters were consistent with Figure 2a, as was Upper Rice Lake bridging the two clusters (Figure 2b), while *Z. aquatica* formed a more distinct cluster in principal coordinate 2. Natural Stand populations from the SCR fell between *Z. aquatica* and the rest of the MN *Z. palustris* populations, with only one *Z. palustris* sample from Dahler Lake overlapping with each of these groups. Other populations appear to cluster along a geographical gradient rather than by watershed. For example, the samples from Clearwater and Ottertail Rivers in the RRN, and Plantagenet and Shell Lakes in the UMR, which are located near one another, also grouped closely together (Figure 2b; Table S2). There was also a more defined population structure among UMR populations with several smaller grouping patterns within the larger two clusters including Necktie River with Garfield Lake, and Bass Lake with Dahler and Decker Lakes populations.

Compared to the Natural Stand collection, there was far less genetic structure evident among genotypes of the 4 cultivars and 10 breeding lines of the Cultivated collection (Figure 2a and c). When the Cultivation collection was analyzed alone, the first and second principal coordinates explained 10% and 7% of the variation, respectively (Figure 2c). The most distinct cluster within the Cultivated collection consisted of samples of ‘K2’, which is a cNWR cultivar released in 1972 that has been kept spatially isolated from other cNWR populations for at least the last 20+ years. While some grouping was present between individuals of a variety or breeding population, there was an overall lack of population structure among Cultivated materials.

The UPGMA clustering analysis (Figure S3) was consistent with some of the clustering observed in the PCoA (Figure 2a-c). Monophyletic clades could be defined for many of the sampling locations, which included the majority of the samples from those populations. Notably, all samples from Clearwater River clustered, as did 98% of the samples from Ottertail River and 92% of the samples from Lake Plantagenet. In total, over 80% of samples within individual populations clustered together except for Upper Rice Lake, Shell Lake and Phantom Lake, which were distributed throughout the tree. Decker Lake and Bass Lake samples each formed two separate clusters distributed through the tree.

Four primary clades were identified in the UPGMA tree. To begin, a *Z. aquatica* clade was identified, which contained the vast majority of the *Z. aquatica* samples but had low bootstrap support (<50%). An additional 25 samples from 8 different Natural Stand populations and 2 Cultivated populations did not cluster with anything else and were found at the base of the tree. The lack of clustering within these samples is likely unknown at this time. The remaining three clades included a clade with samples from Necktie (UMR), Garfield (UMR), and Plantagenet (UMR) Lakes; a clade with samples from Upper Rice (RRN), Phantom (SCR), Dahler (UMR), Decker (UMR) and Bass (UMR) Lakes as well as Cultivated samples; and a clade with samples from Clearwater River (RRN), Shell Lake (UMR) and Ottertail River (RRN).

Within the tree, 91% of Cultivated samples grouped together and formed a cluster with 40% of samples from Upper Rice Lake, 14% each from Phantom and Bass Lakes, 8% from Dahler Lake, 6% each from Shell and Decker Lakes, and a single sample each from Lake Plantagenet and *Z. aquatica* populations. Consistent with the PCoA, there was minimal structure within Cultivated samples. However, unlike the PCoA, samples from Shell Lake, Necktie River, and Lake Plantagenet as well as a single sample of *Z. aquatica* fell within the clade containing Cultivated samples. A second Cultivated cluster at the base of the tree consisted entirely of a subset of K2 samples (10/20).

Using STRUCTURE analyses from *K*=*2*-*14*, evidence of admixture can be seen across species, geographic origin, and germplasm type (Figures 3 and S4), similar to the results of the PCoA and UPGMA analyses. Although there was a significant decrease in the DeltaK statistic between *K=2* and *K=3*, the lowest value was found at *K=5* (Figure S5). Most notably, *Z. aquatica* separated from the Natural Stand populations for the first time at *K=5*, while the Cultivated collection formed its own cluster and the Natural Stands grouped similarly to the PCoA and UPGMA plots (Figure 3). The vast majority of the collections showed limited admixture (<1%) between different populations with the exceptions of Upper Rice Lake and Phantom Lake. Upper Rice Lake showed heavy admixture with Decker Lake, Dahler Lake, Bass Lake, Ottertail River, and Shell Lake populations. Phantom Lake showed an average of 21.43% admixture with the Cultivated materials. Other population groupings were also informative, for example at *K=3*, we observed the Cultivated collection separate into its own cluster, while the other two clusters consisted of the Natural Stand populations and *Z. aquatica* (Figure 3).

**Figure 3.**
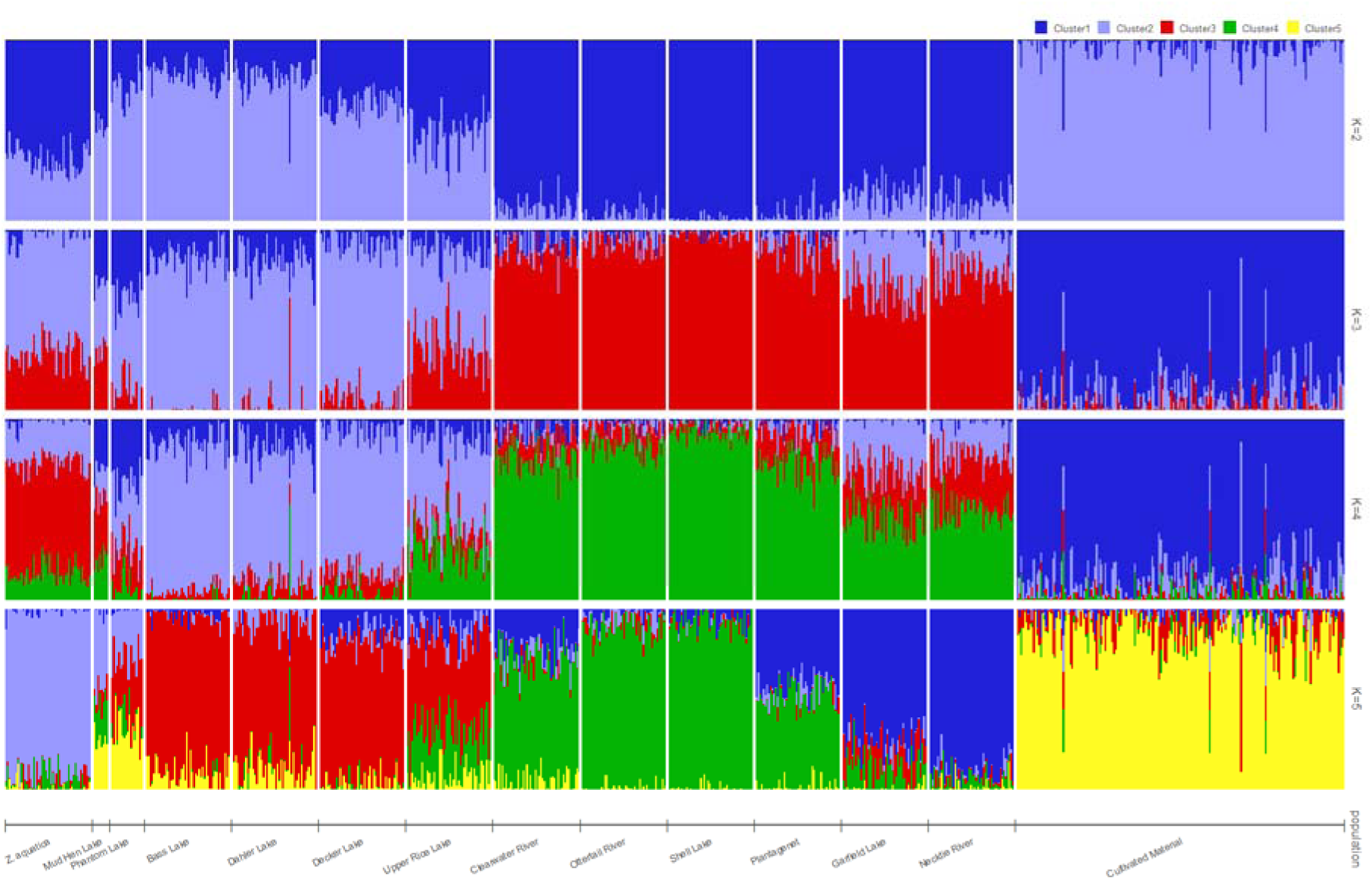
Population structure analysis of Northern Wild Rice (NWR; *Zizania palustris* L.) Natural Stand and Cultivated collections using the program STRUCTURE with 10,000 reps and a burn-in length of 1,000 for *K=2-5*.

Interesting patterns were found at higher *K* values as well. At *K=10*, *Z. aquatica* and the Cultivated collection remained largely unchanged from *K=5* as did Bass Lake, Clearwater River, and Necktie River populations (Figure S4). Populations with high admixture including Phantom and Upper Rice Lakes also remained largely unchanged. However, the Mud Hen Lake population, which showed high admixture at *K=5*, formed a unique cluster at *K=10*, possibly owing to its distance from other sampling sites, as one of only two lakes collected in Wisconsin. Mud Hen Lake was further delineated at *K=14*, splitting into two unique clusters. The results of *K=14* were otherwise consistent with *K=10* (Figure S4).

### Analysis of Molecular Variance

Analysis of Molecular Variance (AMOVA) was conducted between several different groupings including: 1) Natural Stand populations vs. Natural Stand populations; 2) Cultivated lines vs. Cultivated lines; 3) Natural Stand populations vs. Cultivated lines; and 4) *Z. palustris* individuals vs. *Z. aquatica* individuals. The AMOVA results revealed more variation within rather than between groups (Table 2). All the comparative groups identified that 3.37%-8.10% of the variation could be attributed to differences among the groups, rather than within. The highest and lowest variation among groups were identified in the Natural Stand vs Natural Stand analysis (8.10%) and the Cultivated vs Cultivated analysis (3.37%), respectively (Table 2).

**Table 2.** Analysis of Molecular Variance (AMOVA) of a Northern Wild Rice (NWR; *Zizania palustris* L.) diversity collection grouped by germplasm and species type based on 5,955 single nucleotide polymorphism (SNP) markers generated via genotyping-by-sequencing (GBS).

| Grouping | Source of variation | df | MS | Sigma | % of total | Probability |
| --- | --- | --- | --- | --- | --- | --- |
| Natural stand vs. Natural stand | Variation among populations | 12 | 7440.64 | 133.71 | 8.10 | 0.001 |
|  | Variation within populations | 567 | 1516.08 | 1516.08 | 91.90 | 0.001 |
|  | Total of variation | 579 | 1638.87 | 1649.78 | 100.00 | 0.001 |
| Cultivated vs. Cultivated | Variation among populations | 13 | 1592.63 | 18.31 | 1.33 | 0.001 |
|  | Variation within populations | 172 | 1360.34 | 1360.34 | 98.67 | 0.001 |
|  | Total of variation | 185 | 1376.66 | 1378.65 | 100.00 | 0.001 |
| Natural stand vs. Cultivated | Variation among populations | 1 | 20,955.05 | 68.42 | 4.09 | 0.001 |
|  | Variation within populations | 765 | 1604.18 | 1604.17 | 95.91 | 0.001 |
|  | Total of variation | 766 | 1629.44 | 1672.60 | 100.00 | 0.001 |
| *Z. palustris* vs. *Z. aquatica* | Variation among populations | 1 | 9517.98 | 86.37 | 5.05 | 0.001 |
|  | Variation within populations | 578 | 1625.24 | 1625.23 | 94.95 | 0.001 |
|  | Total of variation | 579 | 1638.87 | 1711.61 | 100.00 | 0.001 |

\*df, degrees of freedom; MS, mean of squares

### Gene Flow

*FST* values were calculated to compare the genetic differentiation within and among the Natural Stand and Cultivated collections (Figure 4). Overall, genetic differentiation between the Natural Stand populations was 0.09, with a range of 0.04 - 0.16. Pairwise comparisons that included Mud Hen Lake had the highest *FST* values, most above 0.13, indicating high differentiation from other Natural Stand populations. The Mud Hen Lake population was most similar to Phantom Lake (0.09) and Upper Rice Lake (0.01) populations. Overall, Necktie River and Garfield Lake (*FST* =0.04) and Decker and Upper Rice Lakes (*FST* =0.04) were the most similar populations. As seen with the results of the PCoA, Upper Rice Lake appeared similar to a number of other Natural Stand populations including Bass, Decker, and Phantom Lake populations, which all had *FST* values below 0.05. Although most pairwise comparisons between *Z. aquatica* and the Natural Stand populations had *FST* values above 0.1, Upper Rice Lake and Phantom Lake values were slightly lower at *FST* = 0.08 and 0.07, respectively.

**Figure 4.**
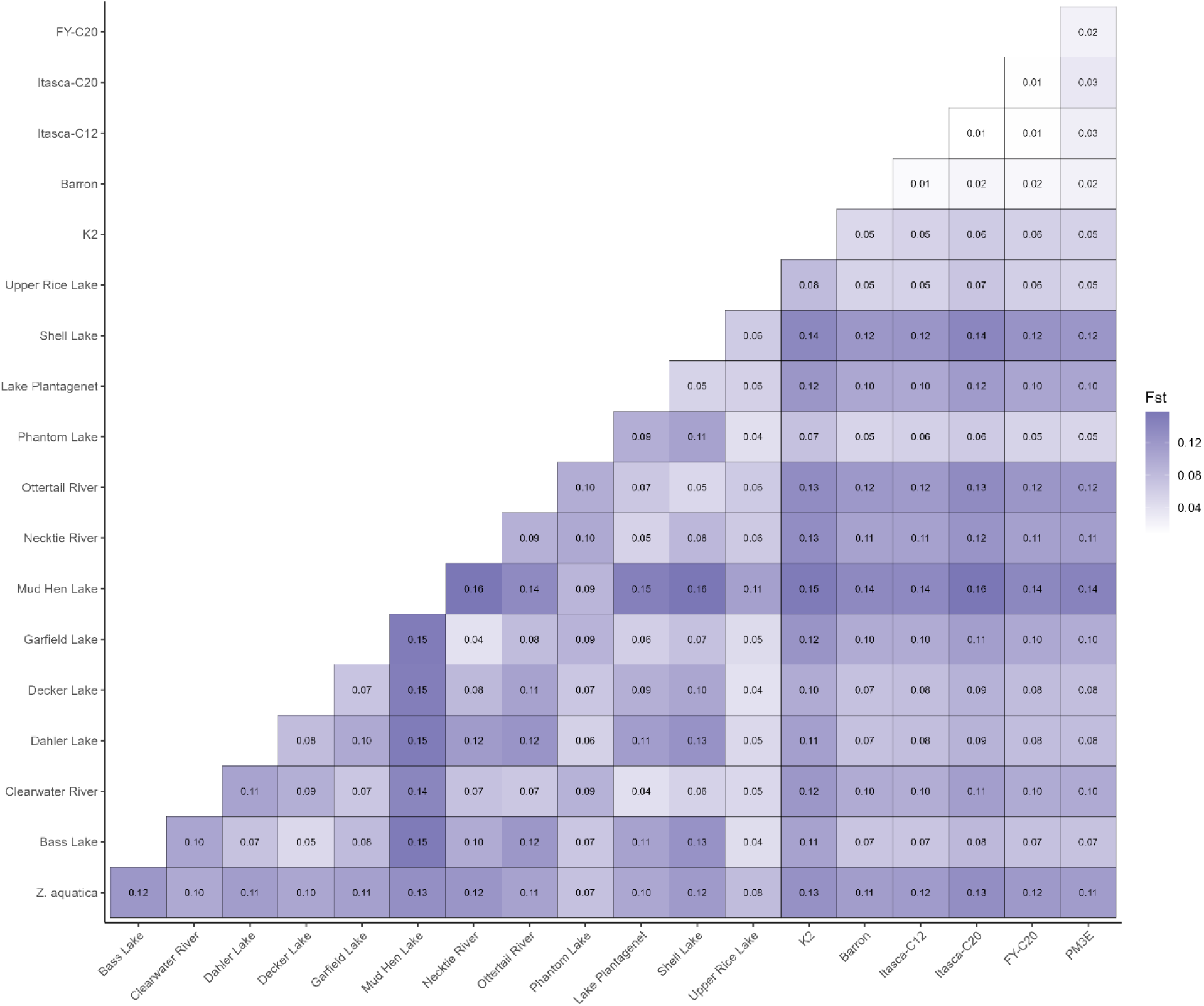
Fixation index (*F_ST_*) values derived using the weighted Weir and Cockerham method (Weir and Cockerham, 1984) for a Natural Stand collection and a Cultivated collection of Northern Wild Rice (NWR; *Zizania palustris* L.).

The pairwise comparisons between Cultivated populations had much lower *FST* values, with an average of 0.03 and a range of 0.01 - 0.06, indicating less differentiation than the Natural Stand populations. For pairwise comparisons between Natural Stand and Cultivated populations, the average *FST* value was 0.10, with a range of 0.05 - 0.16. Phantom lake showed the highest similarity to the Cultivated populations, with *FST* values below 0.06 for all comparisons except that with K2 (*FST*=0.07). Mud Hen Lake comparisons with Cultivated populations had the highest *FST* values, all falling above 0.13, similar to its comparisons with other Natural Stand populations. *Z. aquatica* pairwise comparisons also had relatively high *FST* values, with a range of 0.11 - 0.13.

To further examine the basis for the observed genetic differentiation, we used a Mantel test to look for correlation between genetic and geographic distances of the Natural Stand collection. A positive correlation between the two was observed (r^2^ = 0.4011) and the best fit line yielded a regression equation of y = 0.1 + 0.0002x (Figure S6). Permutation tests showed that the observed results were unlikely to have occurred by chance (*p*-value < 0.05) (Figure S7). We were also interested in whether we could detect admixture between our Natural Stand and Cultivated collections. Using *D*-statistics, we analyzed three groups including one Cultivated group, two Natural Stand groups based on PCoA and STRUCTURE analyses, along with *Z. aquatica*, which served as an outgroup, and found that there was no significant admixture between the groups (Z=1.66, *p*=0.098; Table S6).

### The Temporal Collection

We examined the effect of time on the genetic diversity of two Natural Stand populations collected in 2010 and 2018, using the same approaches used to compare Natural Stand and Cultivated collections. The first principal coordinate of the PCoA for the Temporal collection separated Garfield and Shell Lakes, demonstrating population level differences. Meanwhile, the second coordinate, which explained 7% of the variation, was largely attributed to the temporal variation between Shell Lake samples collected in 2010 vs 2018 (Figure 2d). Within the first two principal components, the Garfield Lake population did not separate by time points. The Shell Lake population, on the other hand, demonstrated a wider range of variation in 2010 than it did in 2018. Additionally, a small unique cluster, not identified in 2010, was identified in 2018 (Figure 2d). Similar patterns were found in the UPGMA tree with monophyletic clades defined for both lakes (Figure S8). There was an even distribution of clustering patterns between Garfield Lake samples collected in 2010 and 2018. For Shell Lake, we observed one unique cluster consisting of 76% of the 2010 samples and 22% of the 2018 samples. The remaining samples showed minimal clustering with few differences between samples collected at different time points (Figure S8). The pairwise comparisons of each lake’s genetic differentiation over time further substantiated these findings as Garfield Lake 2010 vs. 2018 comparison had a lower F*ST* value of 0.0004 and Shell Lake in the same two time periods had a higher value of 0.0125 (Figure 4).

### Scanning for Signatures of Selection

To identify genomic regions potentially subjected to selection or other genetic bottlenecks in cNWR, we calculated genome-wide statistics for nucleotide diversity (π), *FST*, and XP-CLR tests (Figure 5). While we observed considerable similarity between π values of the Natural Stand and Cultivated collections across the genome (Figure 5a), we identified 25 bins with negative Tajima’s D values in the Cultivated collection (Table S7). The most negative Tajima’s D was −0.192 at ~50 Mb on ZPchr0009. Genome-wide scans of *FST* values calculated between Natural Stand and Cultivated collections identified 166 individual pairwise comparisons above 0.10; 63 above 0.20; 24 above 0.30; 9 above 0.40; and 3 above 0.50 (Figure 5b). Genomic positions for these hits are listed in Table S7. The top 1% of *FST* scores were identified on ZPchr0007, ZPchr0009, and ZPchr0013. Finally, we identified 9 regions of the genome with XP-CLR scores above 40 and 4 regions with scores above 60 (Figure 5c). The top 1% of these scores included 468 SNPs, including 141 hits on ZPchr0005 between 8.5-9.7 Mb, 112 hits on ZPchr0006 between 1.2-1.4 Mb, 22 hits on ZPchr0011 between 1.3-1.4 Mb, and 189 hits on ZPchr0013 between 6.2-6.8Mb (Table S7). Allele frequencies of SNPs in these regions were all fixed in the Cultivated collection and ~0.50/0.50 in the Natural Stand collection.

**Figure 5.**
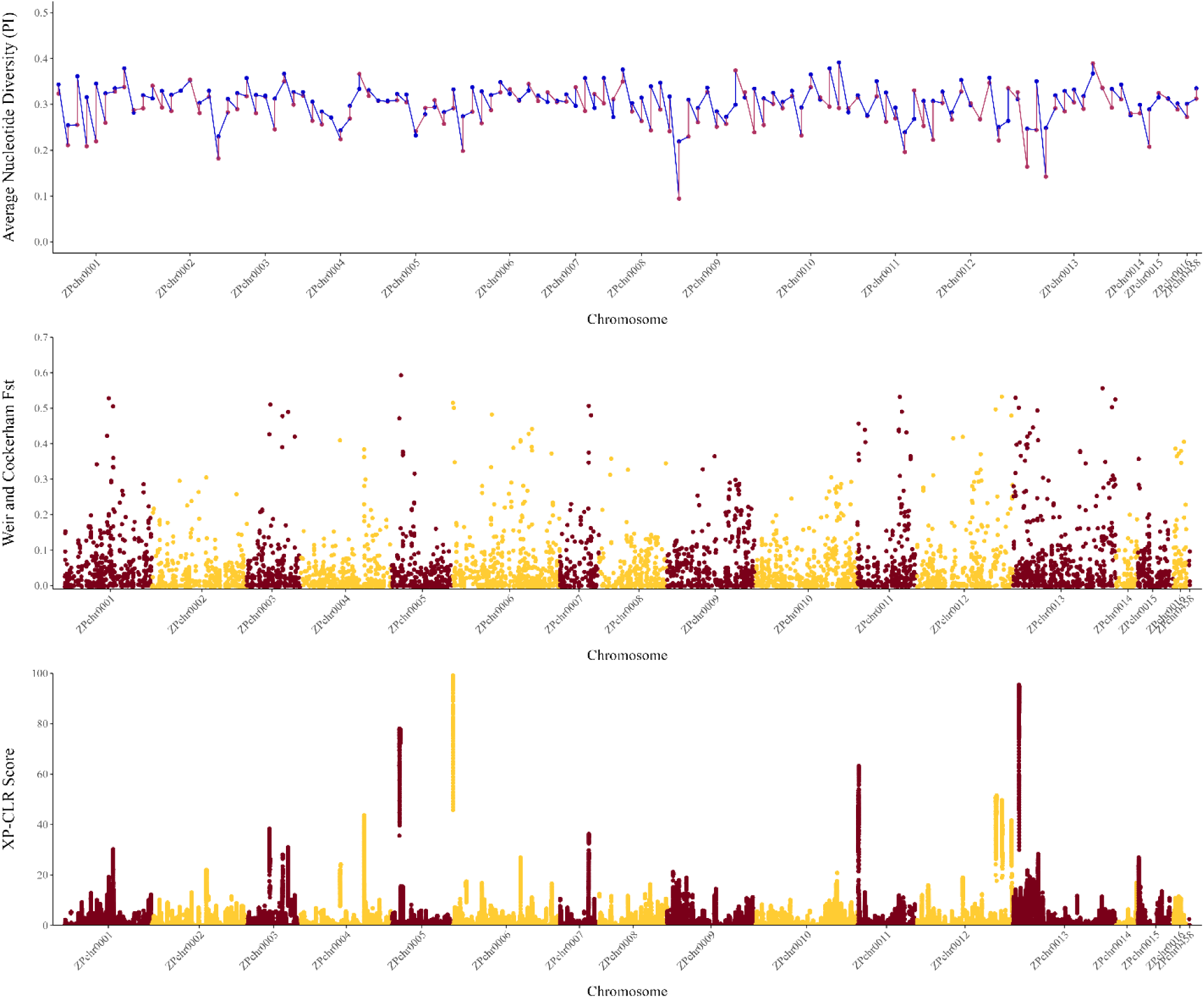
Genome-wide scans of a.) nucleotide diversity (π; Nei, 1987) of a Natural Stand collection (blue) and a Cultivated collection (red) of Northern Wild Rice (NWR; *Zizania palustris* L.); b.) Fixation index (*F_ST_*) values; and c.) XP-CLR scores.

No putative regions of interest were identified across all three tests. However, 11 5-Mb regions with the top 1% of Tajima’s D and *FST* scores were identified, including 2 regions on ZPchr0001; 2 regions on ZPchr0002; 2 on ZPchr0008; 2 regions on ZPchr0009; 1 on ZPchr0010; 1 on ZPchr0012; and 1 on ZPchr0015 (Table S7). Two regions, less than 1 Mb each, with overlapping top 1% of *FST* and Xp-CLR scores were identified on ZPchr0011 and ZPchr0013.

## Discussion

### Assessment of Northern Wild Rice via Genotyping-By-Sequencing

Cost-effective sequencing technologies that are capable of generating robust sets of genome-wide molecular markers, such as GBS, are providing researchers, especially those working with complex plant genomes and limited public resources, an avenue for rapid variant detection ^64–68^. In this study, we developed a large genome-wide SNP dataset, aligned to the NWR reference genome, to assess the relationship between natural and cultivated populations as well as to provide a basis for future breeding and conservation studies. However, we want to emphasize that this GBS approach, which is based on *Btg1* and *TaqI* restriction enzymes, may have introduced a bias in allele frequencies due to polymorphisms in restriction sites ^69^, which could have led to slightly skewed inferences regarding the population genetics of NWR.

### Relationships within the Zizania Genus

Species of *Zizania* are endemic to North America ^70,71^ and split from the Oryzinae subtribe 20-30 million years ago (MYA) ^34,72,73^. Following this split, the Zizaniinae subtribe is hypothesized to have experienced a radiation of speciation across North America and into eastern Asia as individuals made it over the Bering land bridge, leading to the speciation of *Zizania latifolia* ^74^. Comparison of the *Z. palustris* and *Z. latifolia* genomes suggest that the two species split 6-8 MYA ^34^. Extant North American species split 0.7-1.1 MYA; this split was likely precipitated by increases in the habitat range of the *Zizania* progenitor species as climatic conditions shifted over the last million years. ^71,75^. Evidence suggests that *Zizania texana*, an endangered species living in a small stretch of the San Marcos River Valley in the southern US, is a relic, isolated population of the ancestral *Z. palustris* species ^74^. However, the evolutionary relationship between *Z. aquatica*, a species found from the Great Lakes region to the east coast of the continental US, and *Z. palustris* is not well understood. This is likely due to their overlapping range and interspecific crossability ^74,76^. In this study, the UPGMA, STRUCTURE, and principal coordinate analyses showed only moderate support for the separation of *Z. palustris* and *Z. aquatica*. *Fst* values were primarily affected by geographic distance, and highest within *Z. palustris* rather than between *Z. palustris* vs *Z. aquatica* (Figure 4). This may be due to the limited sample size in this study and more research is needed to resolve the complex relationship between these *Zizania* species.

### Structure of Northern Wild Rice Populations is Tied to Geography and a Complex History of Ecosystem Management

Previous genetic analyses of wild populations of NWR have found limited gene flow between populations, and a lack of data to support a correlation between population structure and geographical location ^5,24,74^. In this study, we did not find evidence of significant gene flow among wild populations of NWR, with populations by and large clustering according to their lake or river of origin (Figures 2, 3, and S3). This level of differentiation was also evidenced by the moderate *Fst* values (0.05-0.15)^77^ found between Natural Stand populations (Figure 4). However, the majority of our analyses, including the Mantel test (Figure S6), suggest a geographic basis for population structure in NWR. For example, Garfield Lake and Necktie River, the two closest populations in our study, displayed a high level of similarity with one another (Figure 2b; Figure 3). Therefore, it appears that while gene flow is limited in NWR, it likely occurs between populations in close proximity and more research is needed to understand the spatial dynamics of NWR populations, whose aquatic habitats are often discrete and fragmented. While the primary drivers of gene flow in NWR are not well understood, Lu et al., 2005 found the area and size of a NWR population, along with its degree of isolation, were major factors affecting the genetic variability and gene flow among the NWR populations tested. Additionally, recent pollen travel studies found that most pollen is dispersed within the first 7 m for *Z. palustris* ^78^ and 1.5 m for *Z. texana* ^79^, limiting the likelihood of high levels of gene flow via wind-pollination in the genus.

The historical management and development of lakes across NWR’s natural range have likely contributed to the population structure identified in this study. Efforts to establish new stands of NWR as well as to address declining population sizes have resulted in reseeding efforts across the species’ natural range ^80,81^. For example, Upper Rice Lake (RRN), which is known to have undergone extensive reseeding efforts since the 1930s (Dr. Kimball, *personal communication*), clustered primarily with several UMR populations while showing limited overlap with other RRN populations (Figure 2b; Figure 3). Upper Rice Lake also showed heavy admixture with a number of lakes in STRUCTURE analyses (Figure 3). Taken together, these results suggest that human intervention may have altered the genetic variability and population structure of the Upper Rice Lake population assessed in this study. Additionally, Phantom Lake of the SCR watershed displayed heavy admixture with Cultivated materials. These results were surprising as Phantom Lake is one of the most geographically distant Natural Stand populations from cNWR production in this study, and closer populations displayed little to no admixture with Cultivated materials (Figure 3). However, Phantom Lake is part of the Crex Meadows State Wildlife Area and was artificially created in the 1950s, when a series of levee systems were installed. NWR restoration in this area began in 1991, with 500 lbs (227 kg) of seed sown over the course of three years ^82^. We hypothesize that at least a portion of the seed utilized in these efforts came from cNWR production, further highlighting the complexity of population genetic studies in NWR, as well as the importance of documenting seed sources used in reseeding efforts. We also suggest that future reseeding efforts should not use Phantom Lake populations as a seed source based on the recommendations of the Great Lakes Indian Fish & Wildlife Commission ^83^.

The data presented here for 12 wild populations of NWR likely represents only a small fraction of the species’ genetic diversity. However, even with our small sample size, we were able to identify unique genetic variation within many of the populations in the Natural Stand collection. This indicates that for conservation efforts, it is important to consider populations of NWR individually as they may harbor unique alleles and may be more or less adapted to environmental change. Further studies, using a broader range and more even distribution of sampling locations, will increase our knowledge about the population structure and genetic relationship between wild NWR populations and aid with decision making for future reseeding and other conservation-based efforts.

### Spatio-Temporal Genetic Diversity Analyses Can Aid in Conservation Efforts

Comprehensive monitoring of the spatio-temporal genetic diversity of a species can provide a better understanding of the evolutionary change a species undergoes over time and help to identify targets for conservation efforts. Although present-day genetic diversity studies are available for a wide array of plant species, the majority of spatio-temporal diversity assessments have focused on large agricultural commodity crops ^84–86^. Few studies have focused on natural populations ^87,88^ and many, only theoretically ^89,90^. As NWR is an important target for conservation, monitoring the spatio-temporal diversity of wild populations would provide impactful data for resource managers and environmental agencies interested in the health and preservation of NWR populations across the species natural range.

As a preview of what a more extensive study on the spatio-temporal genetic diversity of NWR could provide, we evaluated two populations, Garfield and Shell Lakes, in 2010 and 2018. Comparing these two time points, we identified a reduction of diversity in samples collected from Shell Lake in 2018 compared to those collected in 2010, while observing limited change in the Garfield Lake population (Figure 2d). This may suggest that Shell Lake has experienced a loss of genetic diversity during the eight years between collection times, while Garfield Lake has not. However, more data is needed to confirm this hypothesis. A wide array of factors could have contributed to this reduction in diversity in Shell Lake, including the 3 - 5 year boom and bust cycles of NWR ^91^, or environmental conditions favorable for specific genotypes in the population’s seed bank. It is also possible that shoreline development and recreation stemming from campgrounds and resorts on Shell Lake could have impacted the health of its native NWR population ^15^.

### Cultivated Northern Wild Rice is Distinct from Natural Stand Populations

Gene flow between domesticated crops and their wild counterparts can have significant impacts on both natural ecosystems and agricultural production systems. Genetic contamination, loss of identity and genetic diversity, and increased weediness are all potential consequences of gene flow ^92^. For these reasons, the extent of gene flow between crops and their wild cohorts has been evaluated in numerous species and found to be dependent on a variety of factors including, but not limited to, mating system (i.e. out-crossing vs selfing), the type and frequency of pollination (i.e. insect vs wind), the selective (dis)advantage of particular domesticated traits (i.e. seed shattering resistance reducing seed dispersal), genetic drift, and genotype × environment interactions ^92–95^. Some studies, such as those in soybean (*Glycine max*), have identified limited gene flow, with domesticated and wild samples separating into monophyletic clades ^96,97^. Other studies have identified significant historical gene flow during domestication, such as Emmer wheat (*Triticum dicoccon*) ^98^, as well as on-going gene flow between crop-weed complexes, such as those in cowpea (*Vigna unguiculata* (L.) Walp) ^99^, pearl millet (*Pennisetum glaucum*)^100^, and species in the *Sorghum* genus ^101,102^.

Given the out-crossing nature of NWR and that cNWR production occurs within the centers of origin and diversity of *Z. palustris*, it is important to understand the extent of gene flow between cultivated and wild populations. This study found that Natural Stand and Cultivated collections are genetically distinct from one another (Figure 2a; Figure 3; Figure S4), indicating minimal gene flow between these two groups and corroborating the results of previous diversity studies in NWR using different marker systems ^5,24,25^. However, based on the 1st principal coordinate from Figure 2a, we identified more similarities between the Cultivated collection and Bass, Decker, and Dahler Lakes than other Natural Stand populations. These lakes are geographically close to the UMN cNWR paddy complex in Grand Rapids, MN and could suggest gene flow. However, it is more likely that this is due to a shared ancestral relationship, as neither STRUCTURE analysis (Figure 3) nor *D*-statistics (Table S6) suggest recent gene flow between the two populations. Importantly, the cultivated germplasm in use today is all descended from natural stand samples originally collected from this geographical region within the UMR watershed. Cultivation and domestication of NWR began in Aitkin, MN and several small enterprises likely gathered seeds from local populations to build their germplasm bases ^13^.

### Domestication and Stewardship of Cultivated Northern Wild Rice

As domestication is a process rather than a specific event, species exhibit varying levels of domestication ^103^. In cereals and other major agricultural crops, seed retention and size, seed dormancy and germination, plant growth habit, and plant size are domestication traits commonly targeted for selection ^104^. The presence of these common traits across multiple taxa is known as the domestication syndrome, which differentiates domesticated species from their wild counterparts. While many of today’s largest agricultural commodity crops have undergone mass selection for thousands of years, the advent of new technologies, such as genomic sequencing, provide today’s plant breeders with new opportunities for the rapid, targeted domestication of new crops ^105^. Additionally, these technologies afford researchers the opportunity to study the domestication process in real-time^106^.

To begin exploring the domestication process of cNWR, we evaluated changes in nucleotide diversity levels and allele frequency distributions between Natural Stand and cNWR populations using Tajima’s D, *FST*, and XP-CLR tests. No significant overlap was identified between the three tests, suggesting there is limited evidence for selective sweeps in cNWR. However, two 1-Mb regions on ZPchr0011 and ZPchr0013 had overlapping top 1% of *FST* and Xp-CLR scores suggesting there is some evidence of genetic changes in cNWR compared to the Natural Stands (Figure 5). A preliminary scan of genes in these two regions identified 5 putative genes whose functions in other species, mainly white rice, are related to drought and salt stresses as well as abscisic acid (ABA) signaling. These included a *60S ribosomal protein kinase 32-like* gene ^107^; a *CBL-interacting protein kinase 32-like* gene ^108^; a E3 ubiquitin-protein ligase RZFP34 isoform X2 ^109,110^; and two copies of *ras-related protein RABC2a* ^111^. Unlike wild populations of NWR, cNWR is grown in man-made irrigated paddies, which are drained shortly after flowering (Principal Phenological Stage 6)^112^ to allow for mechanical harvesting of the grain. Therefore, cNWR experiences conditions similar to upland crops, for which standing water is not available during the development of fruit, ripening, and senescence. These results may suggest that stress-related genes, particularly drought-related genes, were heavily selected for in cNWR germplasm to adapt to this drastic change in environmental conditions compared with its natural habitats. As XP-CLR is more robust than *F* for identifying recent selection events ^63^, we looked at the two additional XP-CLR regions that contained the top 1% of the statistic’s empirical distribution, including a region on ZPchr0005 between 8.5-9.7 Mb and a region on ZPchr0006 between 1.2-1.4 Mb. Within these regions, we identified a *calcium-dependent protein kinase family protein* associated with drought and salt tolerance in white rice ^113,114^; a *2,3-bisphosphoglycerate-independent phosphoglycerate mutase-like* gene involved with chlorophyll synthesis and photosynthesis in white rice ^115^; a *CTD nuclear envelope phosphatase 1 homolog* associated with seed shattering resistance in white rice ^116^; a *KH domain-containing protein SPIN1-like* associated with flowering time in white rice ^117^; and a *pentatricopeptide repeat-containing protein At1g11900 isoform X1* associated with male sterility in *Petunia* ^118^. Two paralogs of *cytochrome P450 714D1-like* were identified on both ZPchr0005 and ZPchr0006 regions of interest. In white rice, this gene is associated with seed dormancy and flowering time ^119,120^. While not in the scope of our current study, we think these regions merit further investigation. Given the significance of NWR to a wide range of stakeholders, it’s important to understand the potential impact of gene flow from cNWR to wild NWR populations. Therefore, while understanding the domestication process in cNWR is important for the plant breeding process, it can also be used to monitor the genetic diversity of natural stands, allowing for better stewardship of these vital populations.

Domestication indices that account for varying levels of domestication have been proposed for several species and typically include: the extent of phenotypic differentiation between the domesticated species and its wild counterparts; the length of a species’ domestication history; whether major genetic changes to the domesticated species have been identified; whether the species has been adapted to agricultural settings through targeted breeding efforts; and the extent of the species’ cultivation ^121–123^. Cultivated NWR is somewhat phenotypically distinct from wild NWR, mainly in its growth habit and seed retention characteristics, which have been made possible through breeding efforts. While the species has a short history of cultivation, its production has expanded to California, which is outside the species’ natural range. For these reasons, we suggest that cNWR should be classified as semi-domesticated.

## Conclusions

Northern Wild Rice is a species with ecological, cultural, and economic importance to the Great Lakes region of North America. Results suggest that wild NWR populations are genetically distinct from each other, and their population structure is influenced by their geographic distribution and possibly, human intervention, such as reseeding efforts. Based on the preliminary temporal data found in this study, we believe it would be beneficial to monitor for shifts in the genetic diversity of NWR populations across both temporal and geographical scales. We also found that wild and cultivated NWR are genetically distinct and that gene flow between the two groups is limited. With the exception of the spatially isolated ‘K2’ breeding population, cNWR germplasm has little population structure and, relative to other commercial crops, appears to be only semi-domesticated. Nevertheless, we found putative selection signals that may be associated with traits that are unique to cultivated NWR including drought tolerance and the bottlebrush panicle type. As the plant breeding process continues, loci with heavy domestication signatures can be used to monitor gene flow between wild and cultivated populations of NWR to expand upon the current conservation and stewardship practices for wild populations.

## Code and Data Availability

All data generated from this project have been deposited at the NCBI Sequence Read Archive under BioProject PRJNA774842. All code for the analyses described can be found at https://github.com/UMNKimballLab/WildRiceGeneticDiversity2022.

## Author Contributions

Lillian McGilp, Matthew Haas, Laura Shannon, and Jennifer Kimball contributed to data analysis, and manuscript preparation and editing. Mingqin Shao managed sequencing. Reneth Millas and Claudia Castell-Miller contributed to data analysis and manuscript preparation. Anthony J. Kern collected leaf tissue samples from natural stands for sequencing and manuscript preparation and editing.

## Supporting Tables

**Table S1.** List of samples in the diversity collection of Northern Wild Rice (NWR; *Zizania palustris* L.) genotyped with 5,955 single nucleotide polymorphism (SNP) markers generated via genotyping-by-sequencing (GBS). HUC 8 watershed designations include Upper Mississippi River (UMR), Red River of the North (RRN), and St. Croix River (SCR) basins. Samples were collected in 2010 and 2018.

| **Sample identification** | **Number of individuals/population** | **Collection Type** | **GPS Coordinates**<br>**Latitud**<br>**Longitud** | **Watershed** |
| --- | --- | --- | --- | --- |
| Bass Lake | 50 | Natural Stand | 47.28716 -<br>93.63132 | UMR |
| Clearwater River | 50 | Natural Stand | 47.51838 -<br>95.46291 | RRN |
| Dahler Lake | 50 | Natural Stand | 46.71888 -<br>93.97281 | UMR |
| Decker Lake | 50 | Natural Stand | 47.63515 -<br>94.40492 | UMR |
| Garfield Lake | 50 | Natural<br>Stand/Temporal | 47.21361 -<br>94.74380 | UMR |
| Mud Hen Lake | 10 | Natural Stand | 45.77005 -<br>92.47173 | SCR |
| Necktie River | 50 | Natural Stand | 47.29363 -<br>94.74724 | UMR |
| Ottertail River | 50 | Natural Stand | 46.38005 -<br>95.86430 | RRN |
| Phantom Lake | 20 | Natural Stand | 45.81839 -<br>92.65074 | SCR |
| Lake Plantagenet | 50 | Natural Stand | 47.36758 -<br>94.91755 | UMR |
| Shell Lake | 50 | Natural<br>Stand/Temporal | 46.94510 -<br>95.48486 | UMR |
| Upper Rice Lake | 50 | Natural Stand | 47.40030 -<br>95.28659 | RRN |
| *Zizania aquatica* | 50 | Natural Stand | 45.94473 -<br>94.24821 | UMR |
| Garfield Lake<br>2010 | 50 | Temporal | 47.21361 -<br>94.74380 | UMR |
| Shell Lake 2010 | 50 | Temporal | 46.94510 -<br>95.48486 | UMR |
| Dwarf | 7 | Cultivated | n/a | n/a |
| Dwarf x PM3E | 3 | Cultivated | n/a | n/a |
| Barron | 25 | Cultivated | n/a | n/a |
| Itasca-C12 | 46 | Cultivated | n/a | n/a |
| Itasca-C20 | 11 | Cultivated | n/a | n/a |
| FY-C20 | 29 | Cultivated | n/a | n/a |
| GP-1 | 3 | Cultivated | n/a | n/a |
| GP-2 | 8 | Cultivated | n/a | n/a |
| K2 | 29 | Cultivated | n/a | n/a |
| K2 x Dwarf x<br>Barron | 9 | Cultivated | n/a | n/a |
| GP-3 | 4 | Cultivated | n/a | n/a |
| PM3E x K2 | 3 | Cultivated | n/a | n/a |
| PM3E | 23 | Cultivated | n/a | n/a |
| Barron x K2 | 6 | Cultivated | n/a | n/a |

**Table S2.**
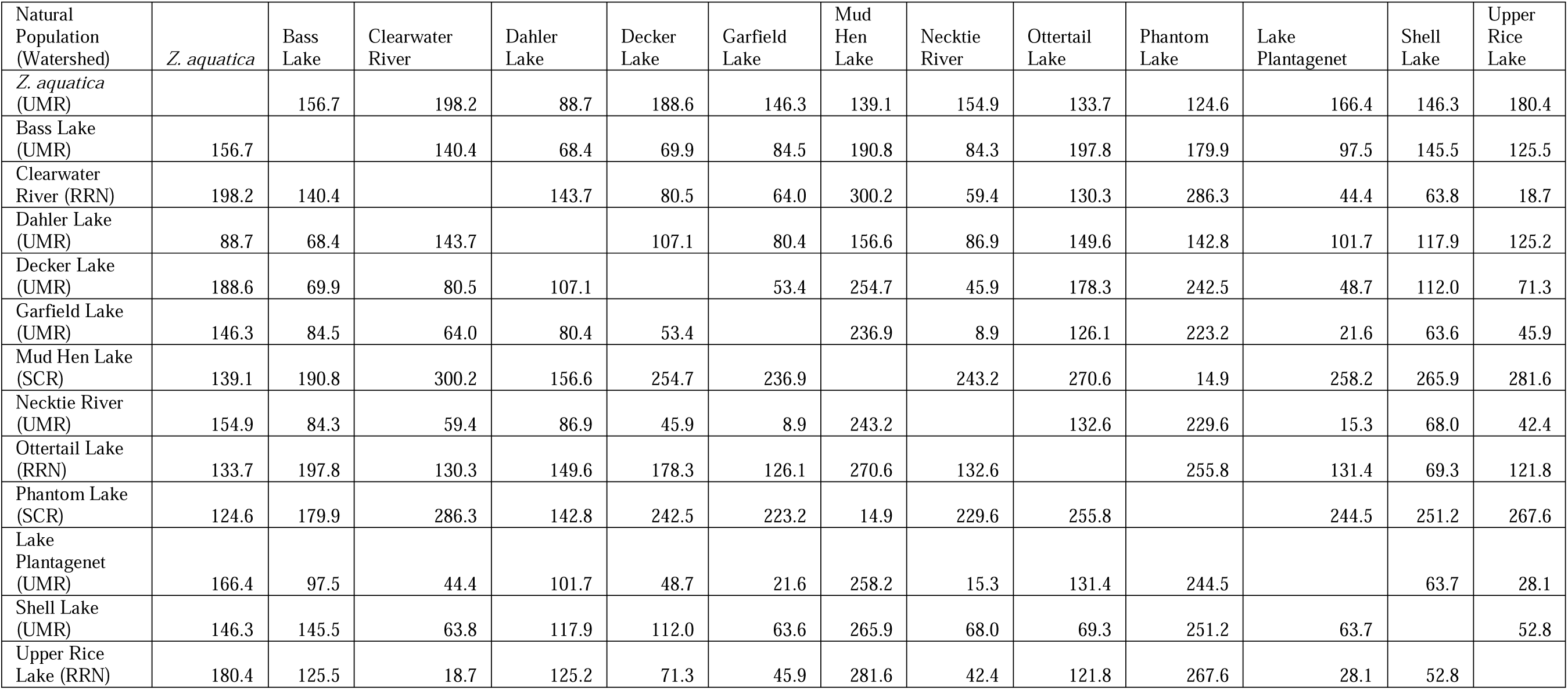
Geographic distance (km) matrix of lakes and rivers where Northern Wild Rice (NWR; *Zizania palustris* L.) and *Zizania aquatica* leaf tissue samples were collected. HUC-8-based watershed designations for the Upper Mississippi River (UMR), Red River of the North (RRN), and St. Croix River (SCR) are included.

| Natural Population (Watershed) | *Z. aquatica* | Bass Lake | Clearwater River | Dahler Lake | Decker Lake | Garfield Lake | Mud Hen Lake | Necktie River | Ottertail Lake | Phantom Lake | Lake Plantagenet | Shell Lake | Upper Rice Lake |
| --- | --- | --- | --- | --- | --- | --- | --- | --- | --- | --- | --- | --- | --- |
| *Z. aquatica* (UMR) |  | 156.7 | 198.2 | 88.7 | 188.6 | 146.3 | 139.1 | 154.9 | 133.7 | 124.6 | 166.4 | 146.3 | 180.4 |
| Bass Lake (UMR) | 156.7 |  | 140.4 | 68.4 | 69.9 | 84.5 | 190.8 | 84.3 | 197.8 | 179.9 | 97.5 | 145.5 | 125.5 |
| Clearwater River (RRN) | 198.2 | 140.4 |  | 143.7 | 80.5 | 64.0 | 300.2 | 59.4 | 130.3 | 286.3 | 44.4 | 63.8 | 18.7 |
| Dahler Lake (UMR) | 88.7 | 68.4 | 143.7 |  | 107.1 | 80.4 | 156.6 | 86.9 | 149.6 | 142.8 | 101.7 | 117.9 | 125.2 |
| Decker Lake (UMR) | 188.6 | 69.9 | 80.5 | 107.1 |  | 53.4 | 254.7 | 45.9 | 178.3 | 242.5 | 48.7 | 112.0 | 71.3 |
| Garfield Lake (UMR) | 146.3 | 84.5 | 64.0 | 80.4 | 53.4 |  | 236.9 | 8.9 | 126.1 | 223.2 | 21.6 | 63.6 | 45.9 |
| Mud Hen Lake (SCR) | 139.1 | 190.8 | 300.2 | 156.6 | 254.7 | 236.9 |  | 243.2 | 270.6 | 14.9 | 258.2 | 265.9 | 281.6 |
| Necktie River (UMR) | 154.9 | 84.3 | 59.4 | 86.9 | 45.9 | 8.9 | 243.2 |  | 132.6 | 229.6 | 15.3 | 68.0 | 42.4 |
| Ottertail Lake (RRN) | 133.7 | 197.8 | 130.3 | 149.6 | 178.3 | 126.1 | 270.6 | 132.6 |  | 255.8 | 131.4 | 69.3 | 121.8 |
| Phantom Lake (SCR) | 124.6 | 179.9 | 286.3 | 142.8 | 242.5 | 223.2 | 14.9 | 229.6 | 255.8 |  | 244.5 | 251.2 | 267.6 |
| Lake Plantagenet (UMR) | 166.4 | 97.5 | 44.4 | 101.7 | 48.7 | 21.6 | 258.2 | 15.3 | 131.4 | 244.5 |  | 63.7 | 28.1 |
| Shell Lake (UMR) | 146.3 | 145.5 | 63.8 | 117.9 | 112.0 | 63.6 | 265.9 | 68.0 | 69.3 | 251.2 | 63.7 |  | 52.8 |
| Upper Rice Lake (RRN) | 180.4 | 125.5 | 18.7 | 125.2 | 71.3 | 45.9 | 281.6 | 42.4 | 121.8 | 267.6 | 28.1 | 52.8 |  |

**Table S3.** List of samples in the diversity collection of Northern Wild Rice (NWR; *Zizania palustris* L.) sorted according to the National Center for Biotechnology Information Short Read Archive (NCBI SRA) BioSample accession numbers. The BioProject ID for the collection is PRJNA774842.

*See External Excel File ‘Table S3’*

**Table S4.** Marker statistics including the-transition/transversion (TsTv) ratios for 5,955 single nucleotide polymorphism (SNP) markers generated via genotyping-by-sequencing (GBS) using the diversity collection of Northern Wild Rice (NWR; *Zizania palustris* L.)

|  | **ZPchr0001** | **ZPchr0002** | **ZPchr0003** | **ZPchr0004** | **ZPchr0005** | **ZPchr0006** | **ZPchr0007** | **ZPchr0008** | **ZPchr0009** |
| --- | --- | --- | --- | --- | --- | --- | --- | --- | --- |
| Scaffold (bp) | 95,470,783 | 103,377,072 | 58,865,324 | 98,769,966 | 66,616,710 | 118,006,097 | 42,614,780 | 75,690,682 | 95,187,837 |
| SNP # | 483 | 435 | 302 | 491 | 350 | 598 | 181 | 367 | 481 |
| SNPs/Mb | 5.06 | 4.21 | 5.13 | 4.97 | 5.25 | 5.07 | 4.25 | 4.85 | 5.05 |
| A/C | 40 | 33 | 30 | 38 | 17 | 60 | 23 | 29 | 44 |
| A/G | 169 | 171 | 108 | 205 | 147 | 197 | 57 | 128 | 181 |
| A/T | 22 | 16 | 12 | 18 | 17 | 26 | 10 | 13 | 16 |
| C/G | 19 | 19 | 17 | 19 | 13 | 34 | 17 | 16 | 17 |
| C/T | 197 | 174 | 109 | 178 | 131 | 233 | 58 | 152 | 196 |
| G/T | 36 | 22 | 26 | 33 | 25 | 48 | 16 | 29 | 27 |
| Ts | 366 | 345 | 217 | 383 | 278 | 430 | 115 | 280 | 377 |
| Tv | 117 | 90 | 85 | 108 | 72 | 168 | 66 | 87 | 104 |
| TsTv ratio | 3.13 | 3.83 | 2.55 | 3.55 | 3.86 | 2.56 | 1.74 | 3.22 | 3.63 |
|  | **ZPchr0010** | **ZPchr0011** | **ZPchr0012** | **ZPchr0013** | **ZPchr0014** | **ZPchr0015** | **ZPchr0016** | **ZPchr0458** | **Genome-wide** |
| Scaffold (bp) | 111,429,323 | 63,217,741 | 105,879,435 | 111,260,184 | 24,058,870 | 39,125,943 | 13,817,767 | 4,333,358 | 1,227,721,872 |
| SNP # | 568 | 285 | 410 | 583 | 91 | 237 | 88 | 5 | 5955 |
| SNPs/Mb | 5.10 | 4.51 | 3.87 | 5.24 | 3.78 | 6.06 | 6.37 | 1.15 | 4.85 |
| A/C | 49 | 21 | 25 | 43 | 7 | 15 | 6 | 0 | 480 |
| A/G | 213 | 109 | 149 | 234 | 30 | 92 | 40 | 2 | 2232 |
| A/T | 25 | 18 | 21 | 23 | 3 | 9 | 1 | 1 | 251 |
| C/G | 31 | 8 | 28 | 28 | 4 | 10 | 4 | 0 | 284 |
| C/T | 210 | 114 | 154 | 212 | 39 | 98 | 31 | 1 | 2287 |
| G/T | 40 | 15 | 33 | 43 | 8 | 13 | 6 | 1 | 421 |
| Ts | 423 | 223 | 303 | 446 | 69 | 190 | 71 | 3 | 4519 |
| Tv | 145 | 62 | 107 | 137 | 22 | 47 | 17 | 2 | 1436 |
| TsTv ratio | 2.92 | 3.60 | 2.83 | 3.26 | 3.14 | 4.04 | 4.18 | 1.50 | 3.15 |

**Table S5.** Polymorphic Information Content (PIC) values for 5,955 single nucleotide polymorphism (SNP) markers generated via genotyping-by-sequencing (GBS) using the diversity collection of Northern Wild Rice (NWR; *Zizania palustris* L.)

|  | Mean | Median | Minimum | Maximum | Standard Deviation |
| --- | --- | --- | --- | --- | --- |
| Barron | 0.1649 | 0.1557 | 0.0216 | 0.3125 | 0.1043 |
| FY-C20 | 0.1639 | 0.1557 | 0.0199 | 0.3125 | 0.1052 |
| Itasca-C12 | 0.1531 | 0.1379 | 0.0166 | 0.3125 | 0.1064 |
| Itasca-C20 | 0.1840 | 0.1829 | 0.0449 | 0.3125 | 0.0981 |
| K2 | 0.1803 | 0.1829 | 0.0261 | 0.3125 | 0.1053 |
| PM3E | 0.1774 | 0.1829 | 0.0292 | 0.3125 | 0.1041 |
| Bass Lake | 0.1484 | 0.1287 | 0.0010 | 0.3125 | 0.1093 |
| Clearwater River | 0.1523 | 0.1369 | 0.0113 | 0.3125 | 0.1084 |
| Dahler Lake | 0.1541 | 0.1373 | 0.0142 | 0.3125 | 0.1079 |
| Decker Lake | 0.1540 | 0.1395 | 0.0091 | 0.3125 | 0.1111 |
| Garfield Lake | 0.1471 | 0.1263 | 0.0091 | 0.3125 | 0.1090 |
| Mud Hen Lake | 0.1924 | 0.1999 | 0.0493 | 0.3125 | 0.0982 |
| Necktie River | 0.1500 | 0.1260 | 0.0113 | 0.3125 | 0.1108 |
| Ottertail River | 0.1528 | 0.1353 | 0.0010 | 0.3125 | 0.1104 |
| Phantom Lake | 0.1637 | 0.1702 | 0.0248 | 0.3125 | 0.1049 |
| (Lake) Plantagenet | 0.1534 | 0.1353 | 0.0010 | 0.3125 | 0.1105 |
| Shell Lake | 0.1477 | 0.1221 | 0.0010 | 0.3125 | 0.1116 |
| Upper Rice Lake | 0.1420 | 0.1141 | 0.0010 | 0.3125 | 0.1088 |
| *Zizania aquatica* | 0.1496 | 0.1313 | 0.0010 | 0.3125 | 0.1107 |

**Table S6.**
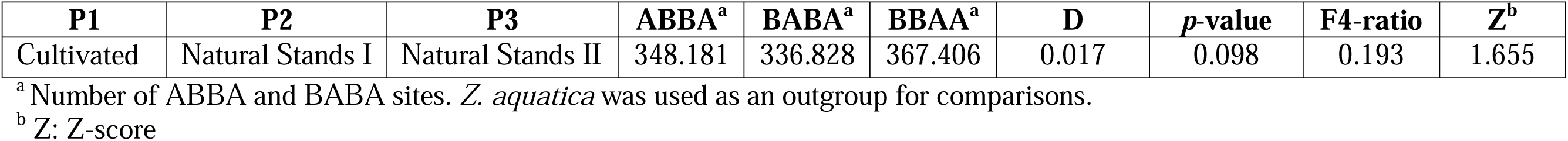
*D-*statistics (ABBA-BABA) results for a diversity collection of Northern Wild Rice (NWR; *Zizania palustris* L.).

**Table S7.** Significant values for Tajima’s D, *F_ST_*, and XP-CLR scores for a diversity collection of Northern Wild Rice (NWR; *Zizania palustris* L.) based on 5,955 single nucleotide polymorphism (SNP) markers generated via genotyping-by-sequencing (GBS).

*See External Excel File ‘Table S7’*

## Supporting Figures

**Figure S1.**
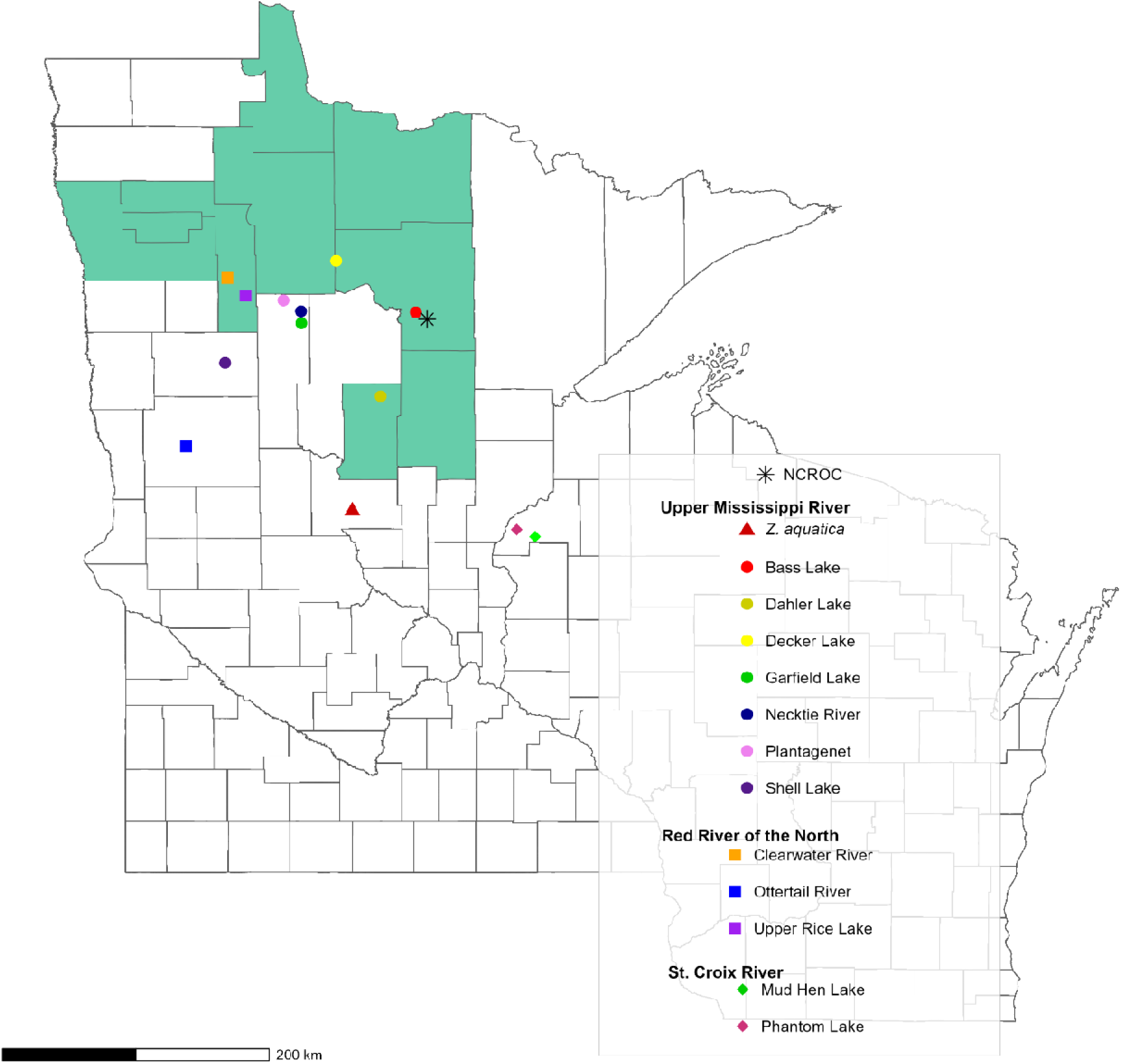
A county-level map of Minnesota and western Wisconsin showing where leaf tissue samples of the Northern Wild Rice (NWR; *Zizania palustris* L.) diversity collection were collected and highlighting counties with significant production of cultivated NWR. Colors and shapes correspond to those featured in the principal component analysis (PCA) plots (Figure 2a-b).

**Figure S2.**
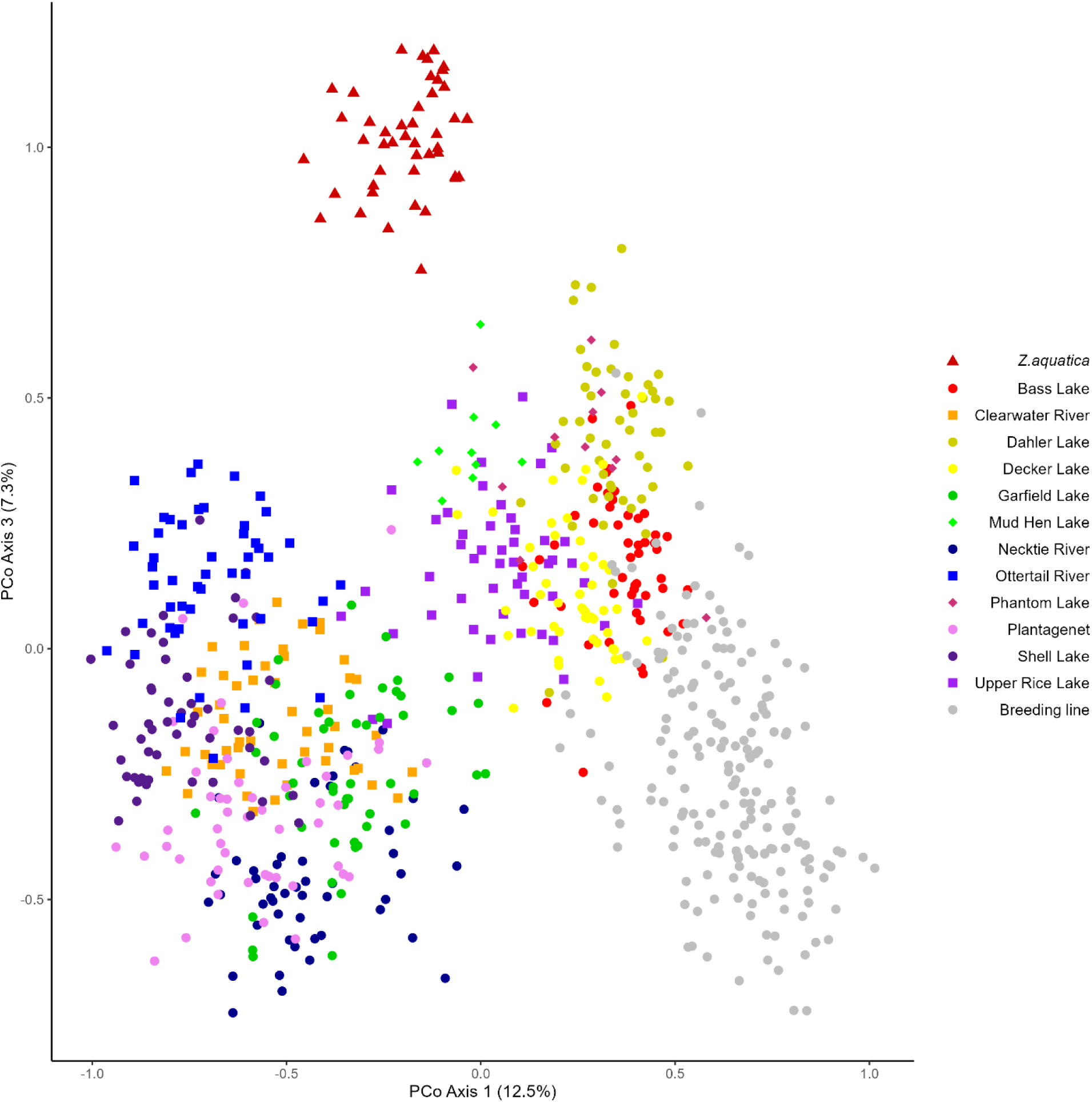
Principal coordinate (PCo) analysis (PCoA) showing the differentiation of the 1^st^ and 3^rd^ PCos of the Natural Stand and Cultivated collections of Northern Wild Rice (NWR; *Zizania palustris* L.).

**Figure S3.**
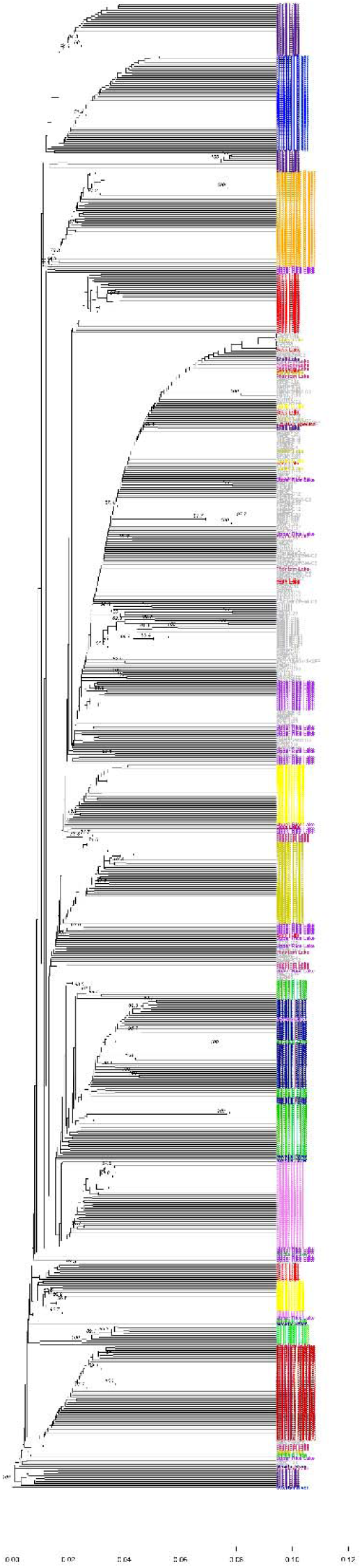
Unweighted pair group method with arithmetic averaging (UPGMA) cluster analysis with bootstrapping of the Natural Stands and Cultivated collections of Northern Wild Rice (NWR; *Zizania palustris* L.).

**Figure S4.**
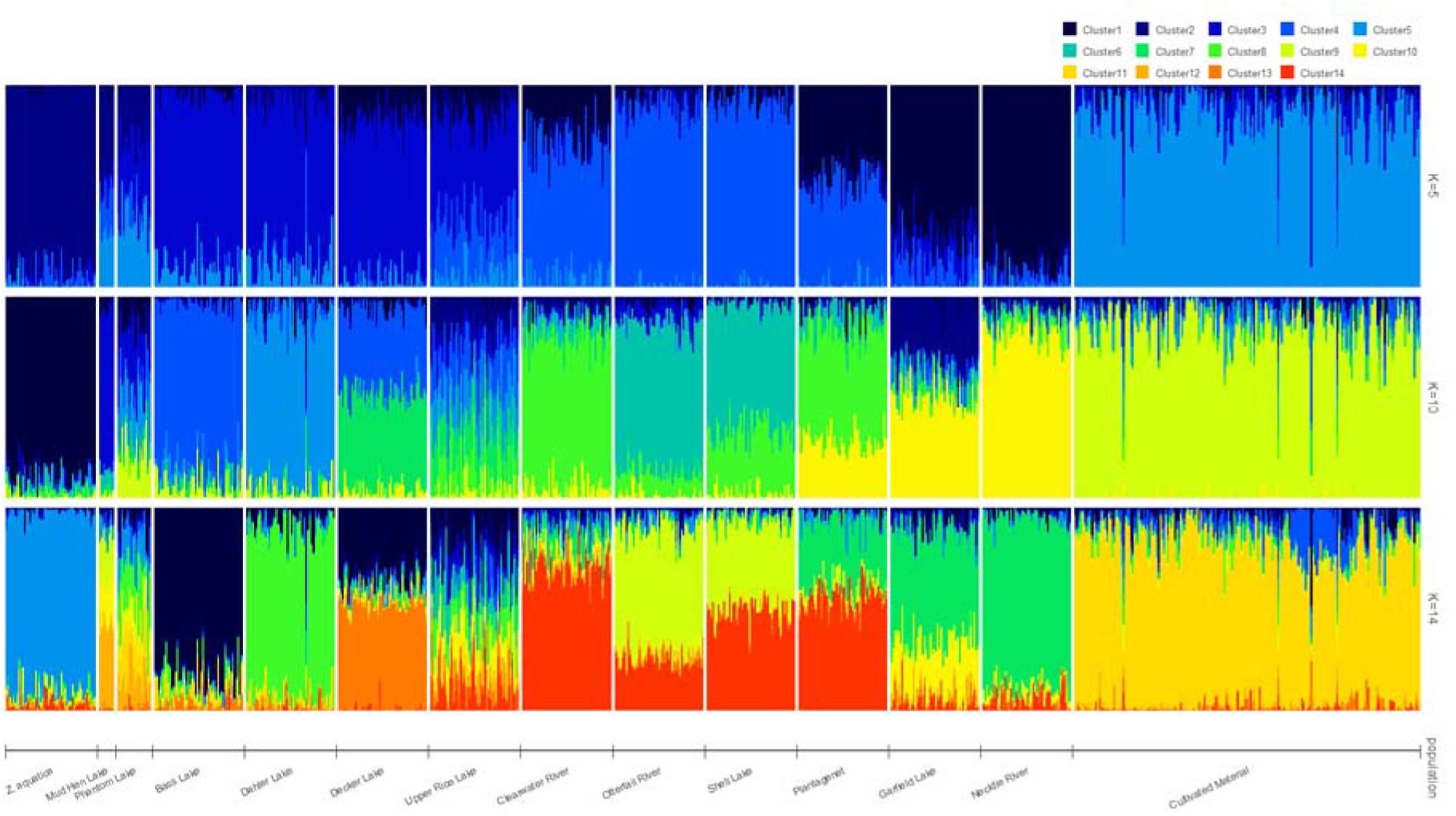
Population structure analysis of Northern Wild Rice (NWR; *Zizania palustris* L.) Natural Stand and Cultivated collections using the program STRUCTURE with 10,000 reps and a burn-in length of 1,000 for *K=5*, *10*, and *14*.

**Figure S5.**
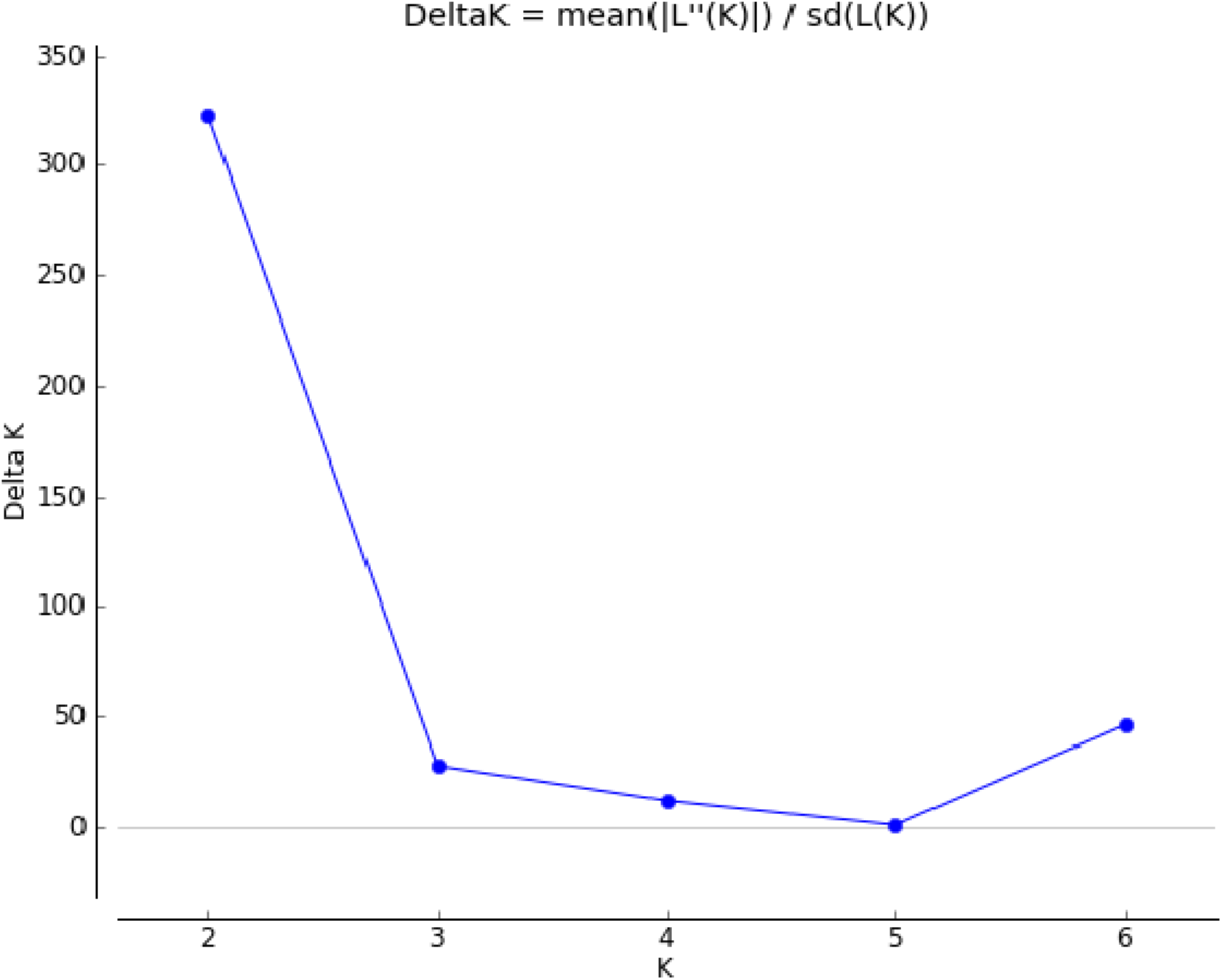
A plot from STRUCTURE HARVESTER performed with the Evanno method based on 5,955 single nucleotide polymorphism (SNP) markers generated via genotyping-by-sequencing (GBS) using the diversity collection of Northern Wild Rice (NWR; *Zizania palustri* L.).

**Figure S6.**
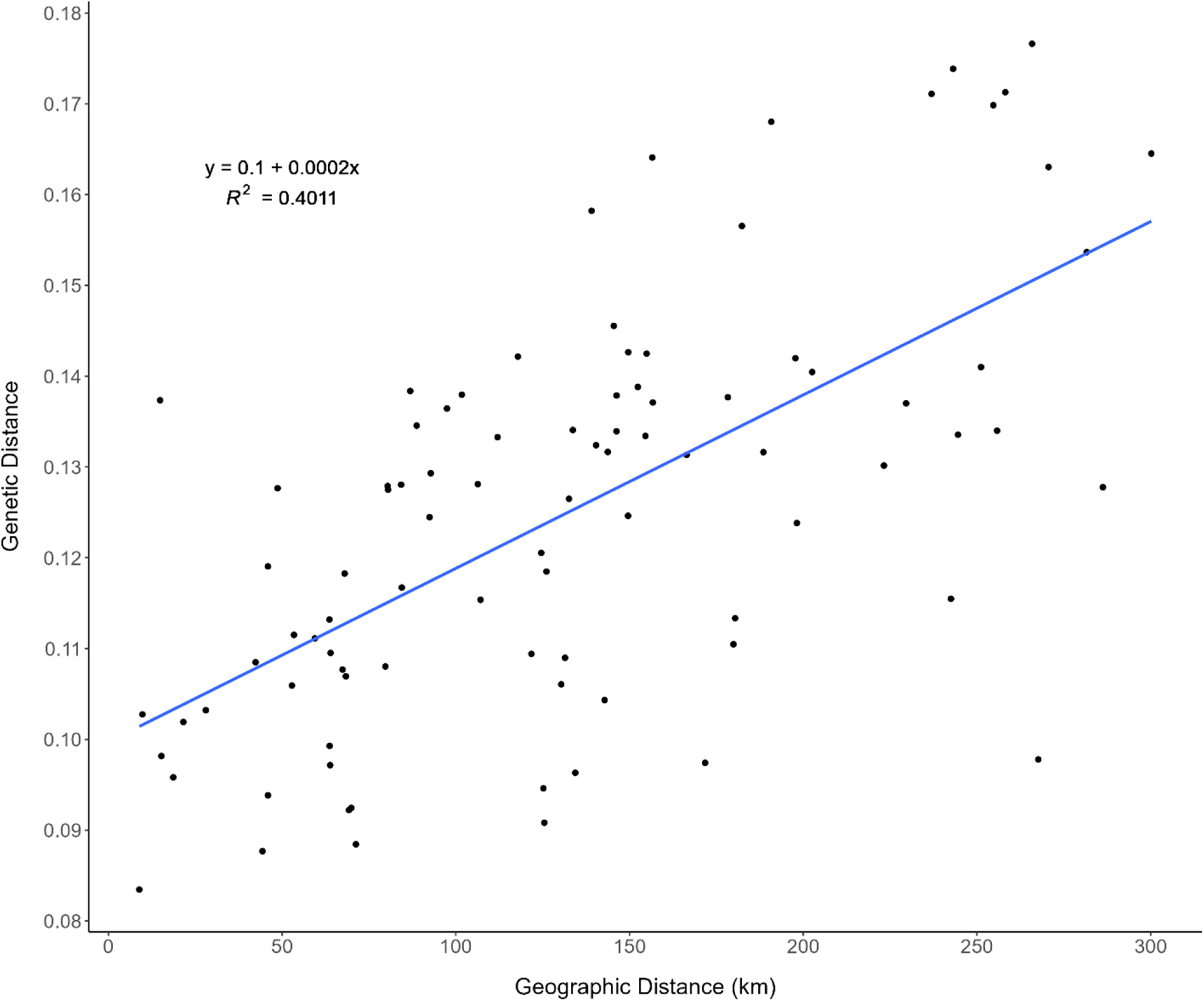
A Mantel test plot showing the correlation between geographic distance (x-axis) and genetic distance (y-axis) for the Natural Stand collection of Northern Wild Rice (NWR; *Zizania palustris* L.). The regression line, y = 0.1 + 0.0002x, was also plotted.

**Figure S7.**
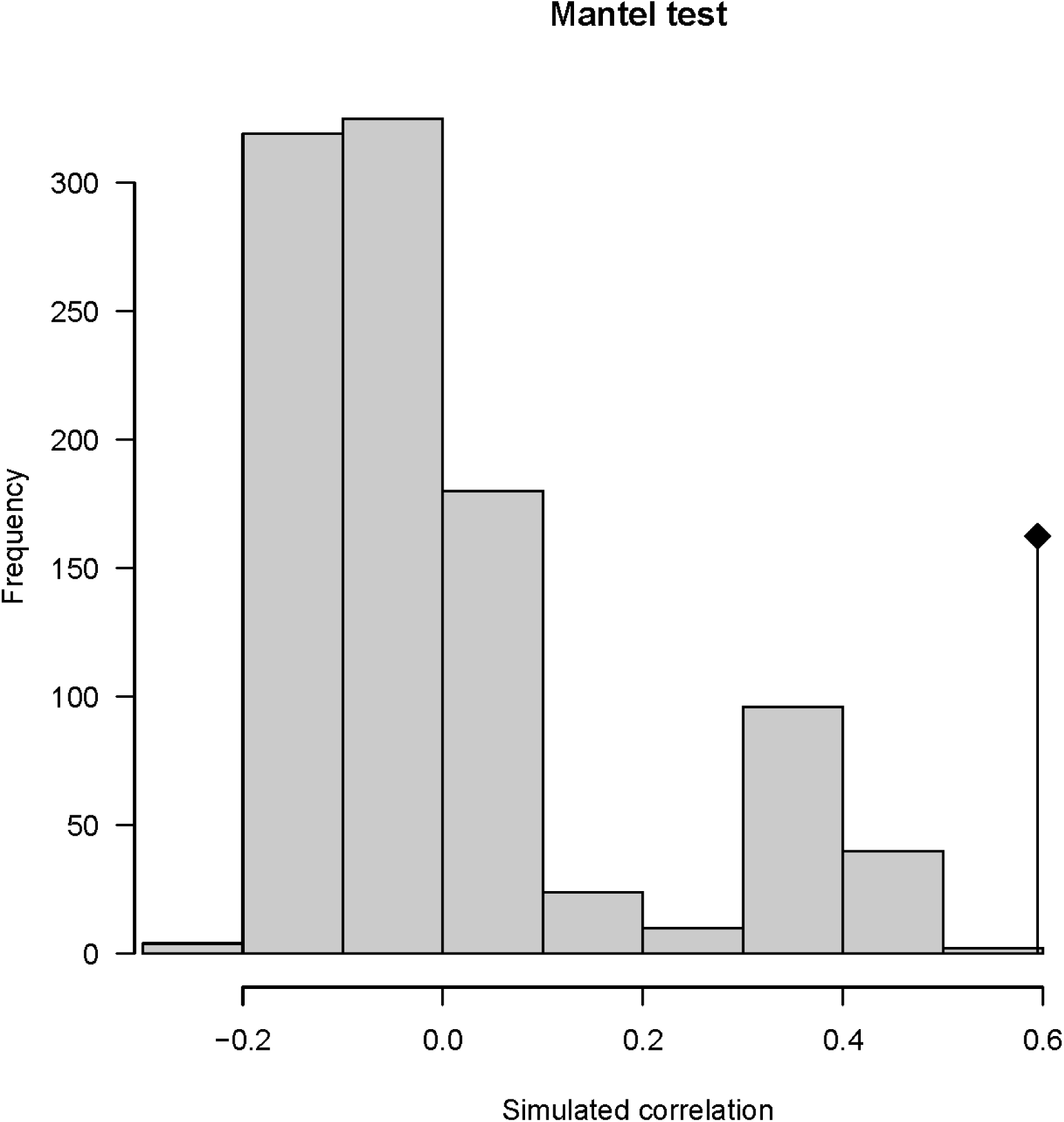
A histogram of the frequency of simulated correlation tests resulting from permutation testing for the Mantel test analysis of the Natural Stand collection of Northern Wild Rice (NWR; *Zizania palustris*). The black diamond with a vertical line beneath it shows the actual correlation value from the Mantel (Figure S6) test using real data. This signifies that results are unlikely to have been reached by chance.

**Figure S8.**
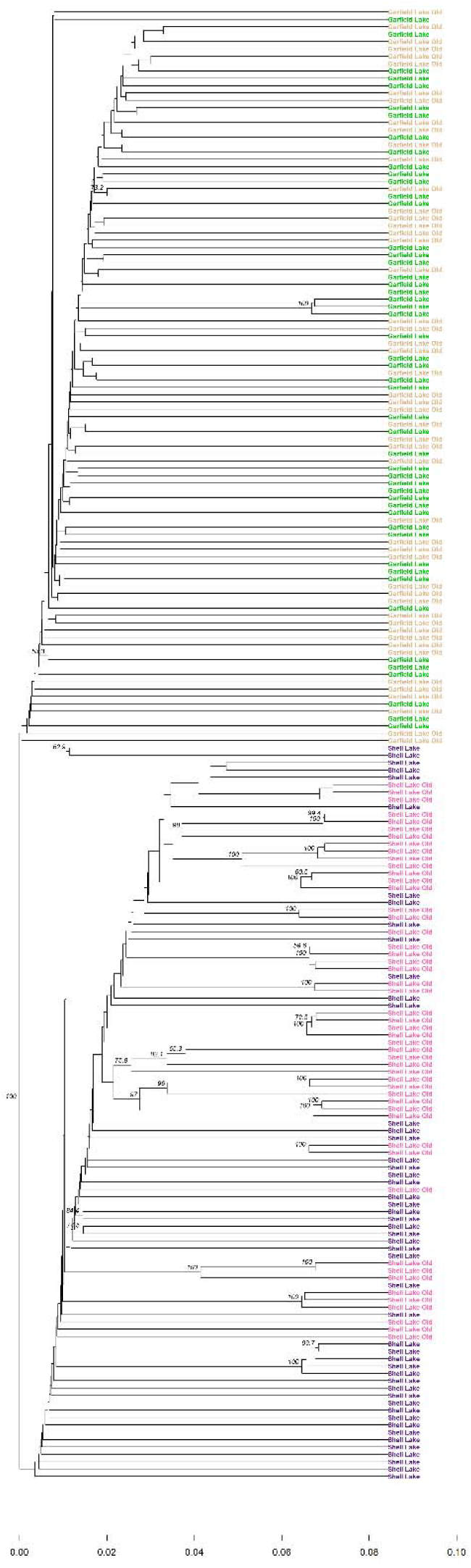
Unweighted pair group method with arithmetic averaging (UPGMA) cluster analysis with bootstrapping of the Temporal collection of Northern Wild Rice (NWR; *Zizania palustris* L.).

## References

1. Elliott, W. A. Wild Rice. in Hybridization of crop plants (eds. Fehr, W. R. & Hadley, H. H.) 721–731 (American Society of Agronomy Madison, WI, 1980).

2. Grombacher, A. W., Porter, R. A. & Everett, L. A. Breeding Wild Rice. in Plant Breeding Reviews 237–265 (John Wiley & Sons, Inc., 1997). doi:10.1002/9780470650073.CH8

3. de Wet, J. M. J. & Oelke, E. A. Domestication of American wild rice (Zizania aquatica L., Gramineae). J. d’agriculture Tradit. Bot. appliquée 25, 67–84 (1978).

4. Drewes, A. D. & Silbernagel, J. Setting up an integrative research approach for sustaining wild rice (Zizania palustris) in the Upper Great Lakes Region of North America. in From Landscape Research to Landscape Planning: Aspects of Integration, Education, and Application (eds. Tress, B., Tress, G., Fry, G. & Opdam, P.) 12, 377–386 (Springer, 2005).

5. Lu, Y., Waller, D. M. & David, P. Genetic variability is correlated with population size and reproduction in American wild-rice (Zizania palustris var. palustris, Poaceae) populations. Am. J. Bot. 92, 990–997 (2005).

6. Myrbo, A. et al. Sulfide Generated by Sulfate Reduction is a Primary Controller of the Occurrence of Wild Rice (*Zizania palustris*) in Shallow Aquatic Ecosystems. J. Geophys. Res. Biogeosciences 122, 2736–2753 (2017).

7. Dean Biesboer, D. Thematic Section: Opinions about Aquatic Ecology in a Changing World. Acta Limnol. Bras. 31, 102 (2019).

8. Desmarais, S. Returning The Rice to the Wild: Revitalizing Wild Rice in the Great Lakes Region Through Indigenous Knowledge Governance and Establishing a Geographical Indication. Lakehead Law J. 3, 36–51 (2019).

9. Moyle, J. B. Wild Rice in Minnesota. J. Wildl. Manage. 8, 177 (1944).

10. Fannucchi, W. Wildlife use of wild rice beds and the impact of rice harvesting on wildlife in east central Minnesota. (University of Wisconsin-Stevens Point, College of Natural Resources, 1983).

11. Rogosin, A. An ecological history of wild rice. (Minnesota Department of Conservation, Division of Game and Fish, 1954).

12. Matson, L., et al. Transforming research and relationships through collaborative tribal-university partnerships on Manoomin (wild rice). Environ. Sci. Policy 115, 108–115 (2021).

13. Oelke, E. A. et al. Wild rice production in Minnesota. Agricultural Extension Service Bulletin 464, 4–5 (1982).

14. Norrgard, R. An overview of threats to the future of wild rice conservation and management. in Manoomin Niikaanisag Wild Rice Coalition Building and Conference 8–11 (2006).

15. Hansen, D. Natural wild rice in Minnesota. A Wild Rice Study Doc. Submitt. to Minnesota Legis. by Minnesota Dep. Resour. (2008).

16. Terrell, E. E., Peterson, P. M., Reveal, J. L. & Duvall, M. R. Taxonomy of north american species of zizania (poaceae). *SIDA*, Contrib. to Bot. 17, 533–549 (1997).

17. Maiz-Tome, L. Zizania aquatica. The IUCN Red List of Threatened Species (2016). Available at: 10.2305/IUCN.UK.2016-1.RLTS.T64326324A67731287.en. (Accessed: 3rd August 2023)

18. Fort, D. J. et al. Toxicity of sulfate and chloride to early life stages of wild rice (*Zizania palustris*). Environ. Toxicol. Chem. 33, 2802–2809 (2014).

19. Probert, R. J. & Longley, P. L. Recalcitrant Seed Storage Physiology in Three Aquatic Grasses (*Zizania palustris*, *Spartina anglica* and *Porteresia coarctata*). Ann. Bot. 63, 53– 63 (1989).

20. Frankel, S. O. H. Genetic conservation of plants useful to man. Biol. Conserv. 2, 162–168 (1970).

21. Ellegren, H. & Galtier, N. Determinants of genetic diversity. Nature Reviews Genetics 17, 422–433 (2016).

22. Sgrò, C. M., Lowe, A. J. & Hoffmann, A. A. Building evolutionary resilience for conserving biodiversity under climate change. Evol. Appl. 4, 326–337 (2011).

23. Santamaría, L. & Méndez, P. F. Evolution in biodiversity policy – current gaps and future needs. Evol. Appl. 5, 202–218 (2012).

24. Kahler, A. L., Kern, A. J., Porter, R. A. & Phillips, R. L. Maintaining food value of wild rice (Zizania palustris L.) using comparative genomics. in Genomics of Plant Genetic Resources 2, 233–248 (Springer Netherlands, 2014).

25. Diller, S. N., McNaught, A. S., Swanson, B. J., Dannenhoffer, J. M. & Ogren, S. Genetic Structure and Morphometric Variation among Fragmented Michigan Wild Rice Populations. Wetlands 38, 793–805 (2018).

26. Collard, B. C. Y., Jahufer, M. Z. Z., Brouwer, J. B. & Pang, E. C. K. An introduction to markers, quantitative trait loci (QTL) mapping and marker-assisted selection for crop improvement: The basic concepts. Euphytica 142, 169–196 (2005).

27. Elshire, R. J. et al. A Robust, Simple Genotyping-by-Sequencing (GBS) Approach for High Diversity Species. PLoS One 6, e19379 (2011).

28. Shao, M., Haas, M., Kern, A. & Kimball, J. Identification of single nucleotide polymorphism markers for population genetic studies in Zizania palustris L. Conserv. Genet. Resour. 12, 451–455 (2019).

29. Streiffer, R. An Ethical Analysis of Ojibway Objections to Genomics and Genetics Research On Wild Rice. Number 12, (2005).

30. Raster, A. & Hill, C. G. The dispute over wild rice: an investigation of treaty agreements and Ojibwe food sovereignty. Agric. Human Values 34, 267–281 (2017).

31. Gross, L. W. The Resolution by the White Earth Anishinaabe Nation to Protect the Inherent Rights of Wild Rice. in Clan and Tribal Perspectives on Social, Economic and Environmental Sustainability 131–140 (Emerald Publishing Limited, 2021).

32. Andrews, S. FastQC: A quality control tool for high throughput sequence data. (2010).

33. Martin, M. Cutadapt removes adapter sequences from high-throughput sequencing reads. EMBnet.journal 17, 10 (2011).

34. Haas, M. et al. Whole Genome Assembly and Annotation of Northern Wild Rice, Zizania palustris L., Supports a Whole Genome Duplication in the Zizania Genus. Zizania Genus 2, 3 (2021).

35. Li, H. Aligning sequence reads, clone sequences and assembly contigs with BWA-MEM. (2013).

36. Li, H. A statistical framework for SNP calling, mutation discovery, association mapping and population genetical parameter estimation from sequencing data. Bioinformatics 27, 2987–2993 (2011).

37. Li, H. et al. The Sequence Alignment/Map format and SAMtools. Bioinformatics 25, 2078–2079 (2009).

38. Danecek, P. et al. The variant call format and VCFtools. Bioinformatics 27, 2156–2158 (2011).

39. Granato, I. S. C. et al. snpReady: a tool to assist breeders in genomic analysis. Mol. Breed. 38, 1–7 (2018).

40. R Core Team. R: A language and environment for statistical computing. (2021).

41. Knaus, B. J. & Grünwald, N. J. <scp>vcfr</scp> : a package to manipulate and visualize variant call format data in R. Mol. Ecol. Resour. 17, 44–53 (2017).

42. Purcell, S. et al. PLINK: A tool set for whole-genome association and population-based linkage analyses. Am. J. Hum. Genet. 81, 559–575 (2007).

43. Oksanen, J. et al. vegan: Community ecology package. (2022).

44. Wickham, H. ggplot2: Elegant graphics for data analysis. (2016).

45. Jombart, T. adegenet: a R package for the multivariate analysis of genetic markers. Bioinformatics 24, 1403–1405 (2008).

46. Jombart, T. & Ahmed, I. adegenet 1.3-1: new tools for the analysis of genome-wide SNP data. Bioinformatics 27, 3070–3071 (2011).

47. Paradis, E. & Schliep, K. ape 5.0: an environment for modern phylogenetics and evolutionary analyses in R. Bioinformatics 35, 526–528 (2019).

48. Kamvar, Z. N., Tabima, J. F. & Gr unwald, N. J. Poppr: An R package for genetic analysis of populations with clonal, partially clonal, and/or sexual reproduction. PeerJ 2014, 1–14 (2014).

49. Kamvar, Z. N., Brooks, J. C. & GrÃ¼nwald, N. J. Novel R tools for analysis of genome-wide population genetic data with emphasis on clonality. Front. Genet. 6, 208 (2015).

50. Jacquemyn, H., Honnay, O., Van Looy, K. & Breyne, P. Spatiotemporal structure of genetic variation of a spreading plant metapopulation on dynamic riverbanks along the Meuse River. Heredity (Edinb). 96, 471–478 (2006).

51. Pritchard, J. K., Stephens, M. & Donnelly, P. Inference of Population Structure Using Multilocus Genotype Data. Genetics 155, 945–959 (2000).

52. Francis, R. M. pophelper: an R package and web app to analyse and visualise population structure. Mol. Ecol. Resour. 17, 27–32 (2017).

53. Earl, D. A. & vonHoldt, B. M. STRUCTURE HARVESTER: A website and program for visualizing STRUCTURE output and implementing the Evanno method. Conserv. Genet. Resour. 4, 359–361 (2012).

54. Evanno, G., Regnaut, S. & Goudet, J. Detecting the number of clusters of individuals using the software structure: a simulation study. Mol. Ecol. 14, 2611–2620 (2005).

55. Excoffier, L., Smouse, P. E. & Quattro, J. M. Analysis of molecular variance inferred from metric distances among DNA haplotypes: application to human mitochondrial DNA restriction data. Genetics 131, 479–491 (1992).

56. Dray, S. & Dufour, A. B. The ade4 package: Implementing the duality diagram for ecologists. J. Stat. Softw. 22, 1–20 (2007).

57. Hijmans, R. geosphere: Spherical trigonometry. (2022).

58. Weir, B. S. & Cockerham, C. C. Estimating F-Statistics for the Analysis of Population Structure. Evolution (N. Y). 38, 1358–1370 (1984).

59. Pembleton, W., Cogan, N. O. I. & Forster, J. W. StAMPP: an R package for calculation of genetic differentiation and structure of mixed-ploidy level populations. Mol. Ecol. Resour. 13, 946–952 (2013).

60. Malinsky, M., Matschiner, M. & Svardal, H. Dsuite Fast *D* statistics and related admixture evidence from VCF files. Mol. Ecol. Resour. 21, 584–595 (2021).

61. Nei, M. Molecular Evolutionary Genetics. Columbia University Press (Columbia University Press, 1987). doi:doi:10.7312/nei-92038

62. Tajima, F. Statistical method for testing the neutral mutation hypothesis by DNA polymorphism. Genetics 123, 585–595 (1989).

63. Chen, H., Patterson, N. & Reich, D. Population differentiation as a test for selective sweeps. Genome Res. 20, 393–402 (2010).

64. Delfini, J. et al. Population structure, genetic diversity and genomic selection signatures among a Brazilian common bean germplasm. Sci. Rep. 11, 2964 (2021).

65. Tuttle, H. K., Del Rio, A. H., Bamberg, J. & Shannon, L. M. Potato soup: analysis of cultivated potato gene bank populations reveals high diversity and little structure. Front. Plant Sci. 15, 1429279 (2024).

66. Migicovsky, Z. et al. Genomic consequences of apple improvement. Hortic. Res. 8, 9 (2021).

67. Bredeson, J. V et al. Sequencing wild and cultivated cassava and related species reveals extensive interspecific hybridization and genetic diversity. Nat. Biotechnol. 34, 562–570 (2016).

68. García-Abadillo, J. et al. Dissecting the complex genetic basis of pre- and post-harvest traits in Vitis vinifera L. using genome-wide association studies. Hortic. Res. 11, uhad283 (2024).

69. Arnold, B., Corbett-Detig, R. B., Hartl, D. & Bomblies, K. RADseq underestimates diversity and introduces genealogical biases due to nonrandom haplotype sampling. Mol. Ecol. 22, 3179–3190 (2013).

70. Xu, X., Ke, W., Yu, X., Wen, J. & Ge, S. Preliminary study on population genetic structure and phylogeography of the wild and cultivated *Zizania latifolia* (Poaceae) based on Adh1a sequences. Theor. Appl. Genet. 116, 835–843 (2008).

71. Xu, X. et al. Phylogeny and biogeography of the eastern Asian–North American disjunct wild-rice genus (*Zizania* L., Poaceae). Mol. Phylogenet. Evol. 55, 1008–1017 (2010).

72. Guo, L. et al. A host plant genome (Zizania latifolia) after a century-long endophyte infection. Plant J. 83, 600–609 (2015).

73. Tang, L. et al. Phylogeny and biogeography of the rice tribe (Oryzeae): Evidence from combined analysis of 20 chloroplast fragments. Mol. Phylogenet. Evol. 54, 266–277 (2010).

74. Xu, X. W. et al. Comparative phylogeography of the wild-rice genus Zizania (Poaceae) in eastern Asia and north America. Am. J. Bot. 102, 239–247 (2015).

75. Walker, S. A. DNA sequence diversity in North American Zizania species. Purdue University ProQuest Dissertations Publishing (ProQuest Dissertations Publishing, 2011).

76. Duvall, M. R. & Biesboer, D. D. Anatomical distinctions between the pistillate spikelets of the species of wild-rice (Zizania poaceae). Am. J. Bot. 75, 157–159 (1988).

77. Wright, S. The Interpretation of Population Structure by F-Statistics with Special Regard to Systems of Mating. Evolution (N. Y). 19, 395–420 (1965).

78. Gietzel, C., Duquette, J., McGilp, L. & Kimball, J. Recessive male floret color for tracking gene flow in cultivated northern wild rice (Zizania palustris L.). Crop Sci. 62, 157–166 (2022).

79. Oxley, F. M., Echlin, A., Power, P., Tolley-Jordan, L. & Alexander, M. L. Travel of pollen in experimental raceways in the endangered texas wild rice (Zizania texana). Southwest. Nat. 53, 169–174 (2008).

80. Brandes, H. Like Gold to Us: Native American Nations Struggle to Protect Wild Rice. Sierra: The Magazine of the Sierra Club (2019). Available at: https://www.sierraclub.org/sierra/gold-us-native-american-nations-struggle-protect-wild-rice.

81. Porter, R. Wildrice (Zizania L.) in North America: Genetic resources, conservation, and use. in North American Crop Wild Relatives: Important Species 2, 83–97 (Springer International Publishing, 2019).

82. Thompson, A. & Luthin, C. S. Wild Rice Community Restoration. in Wetland Restoration Handbook for Wisconsin Landowners 117–122 (Bureau of Science Services - Wisconsin DNR, 2010).

83. David, P., David, L., Stark, H. K., Fahrlander, S. N. A. & Schlender, J. M. Manoomin, Version 1.0. Gt. Lakes Indian Fish Wildl. Comm. [Preprint]. (Accessed December 3, 2020) (2019).

84. Le Clerc, V. et al. Indicators to assess temporal genetic diversity in the French Catalogue: no losses for maize and peas. Theor. Appl. Genet. 113, 1197–1209 (2006).

85. Deu, M. et al. Spatio-temporal dynamics of genetic diversity in Sorghum bicolor in Niger. Theor. Appl. Genet. 120, 1301–1313 (2010).

86. Arya, L., Verma, M., Singh, S. & Verma, R. Spatio-temporal genetic diversity in Indian barley (Hordeum vulgare L.) varieties based on SSR markers. Indian J. Exp. Biol. 57, 545–552 (2019).

87. Ozbek, O., Millet, E., Anikster, Y., Arslan, O. & Feldman, M. Spatio-temporal genetic variation in populations of wild emmer wheat, Triticum turgidum ssp. dicoccoides, as revealed by AFLP analysis. Theor. Appl. Genet. 115, 19–26 (2007).

88. Fačkovcová, Z. et al. Spatio-temporal formation of the genetic diversity in the Mediterranean dwelling lichen during the Neogene and Quaternary epochs. Mol. Phylogenet. Evol. 144, 106704 (2020).

89. Gray, A. Genetic diversity and its conservation in natural populations of plants. Biodivers. Lett. 3, 71–80 (1996).

90. González-Megías, A., Gómez, J. M. & Sánchez-Piñero, F. Spatio-temporal change in the relationship between habitat heterogeneity and species diversity. Acta Oecologica 37, 179–186 (2011).

91. Waheed, A. Manoomin (wild rice) and environmental change at a significant river system of the Lac du Flambeau Band of Lake Superior Chippewa. University of Minnesota Digital Conservancy (University of Minnesota, 2021).

92. Gepts, P. & Papa, R. Possible effects of (trans)gene flow from crops on the genetic diversity from landraces and wild relatives. Environ. Biosaf. Res 2, 89–103 (2003).

93. Sonnante, G., Stockton, T., Nodari, R. O., Becerra Velásquez, V. L. & Gepts, P. Evolution of genetic diversity during the domestication of common-bean (Phaseolus vulgaris L.). Theor. Appl. Genet. 89, 629–635 (1994).

94. Evans, M. M. S. & Kermicle, J. L. Teosinte crossing barrier1, a locus governing hybridization of teosinte with maize. Theor. Appl. Genet. 103, 259–265 (2001).

95. Gepts, P. A Comparison between Crop Domestication, Classical Plant Breeding, and Genetic Engineering. Crop Sci. 42, 1780–1790 (2002).

96. Li, Y.-H. et al. Genetic diversity in domesticated soybean (Glycine max) and its wild progenitor (Glycine soja) for simple sequence repeat and single-nucleotide polymorphism loci. New Phytol. 188, 242–253 (2010).

97. Jeong, S.-C. et al. Genetic diversity patterns and domestication origin of soybean. 132, 1179–1193 (2019).

98. Luo, M. C. et al. The structure of wild and domesticated emmer wheat populations, gene flow between them, and the site of emmer domestication. Theor. Appl. Genet. 114, 947– 959 (2007).

99. Coulibaly, S., Pasquet, R. S., Papa, R. & Gepts, P. AFLP analysis of the phenetic organization and genetic diversity of Vigna unguiculata L. Walp. reveals extensive gene flow between wild and domesticated types. Theor. Appl. Genet. 104, 358–366 (2002).

100. Mariac, C. et al. Genetic diversity and gene flow among pearl millet crop/weed complex: A case study. Theor. Appl. Genet. 113, 1003–1014 (2006).

101. Arriola, P. E. & Ellstrand, N. C. Crop-to-weed gene flow in the genus *Sorghum* (Poaceae): Spontaneous interspecific hybridization between johnsongrass, *Sorghum halepense*, and crop sorghum, *S. bicolor*. Am. J. Bot. 83, 1153–1159 (1996).

102. Sagnard, F. et al. Genetic diversity, structure, gene flow and evolutionary relationships within the Sorghum bicolor wild-weedy-crop complex in a western African region. Theor. Appl. Genet. 123, 1231–1246 (2011).

103. Zeder, M. A., Emshwiller, E., Smith, B. D. & Bradley, D. G. Documenting domestication: The intersection of genetics and archaeology. Trends in Genetics 22, 139–155 (2006).

104. Stalker, H. T., Warburton, M. L. & R, H. J. The Dynamics of Domestication. in Harlan’s Crops and Man (eds. Stalker, H. T., Warburton, M. L. & R, H. J.) 147–170 (American Society of Agronomy, Inc. and Crop Science Society of America, Inc., 2021). 10.1002/9780891186342.ch6

105. Zhang, C. et al. Genome design of hybrid potato. Cell 184, 3873–3883.e12 (2021).

106. Ekar, J. M. et al. Domestication in Real Time: The Curious Case of a Trigenomic Sunflower Population. Agronomy 9, (2019).

107. Ji, W. et al. Quantitative proteomics reveals an important role of GsCBRLK in salt stress response of soybean. Plant Soil 402, 159–178 (2016).

108. Hu, D. et al. Overexpression of MdSOS2L1, a CIPK protein kinase, increases the antioxidant metabolites to enhance salt tolerance in apple and tomato. Physiol. Plant. 156, 201–214 (2016).

109. Ding, S., Zhang, B. & Qin, F. Arabidopsis RZFP34/CHYR1, a Ubiquitin E3 Ligase, Regulates Stomatal Movement and Drought Tolerance via SnRK2.6-Mediated Phosphorylation. Plant Cell 27, 3228–3244 (2015).

110. Shu, K. & Yang, W. E3 Ubiquitin Ligases: Ubiquitous Actors in Plant Development and Abiotic Stress Responses. Plant Cell Physiol. 58, 1461–1476 (2017).

111. Mérida-García, R. et al. High Resolution Melting and Insertion Site-Based Polymorphism Markers for Wheat Variability Analysis and Candidate Genes Selection at Drought and Heat MQTL Loci. Agron. 2020, Vol. 10, Page 1294 10, 1294 (2020).

112. Duquette, J. & Kimball, J. A. Phenological stages of cultivated northern wild rice according to the BBCH scale. Ann. Appl. Biol. 176, 350–356 (2020).

113. Campo, S. et al. Overexpression of a Calcium-Dependent Protein Kinase Confers Salt and Drought Tolerance in Rice by Preventing Membrane Lipid Peroxidation. Plant Physiol. 165, 688–704 (2014).

114. Wei, S. et al. A rice calcium-dependent protein kinase OsCPK9 positively regulates drought stress tolerance and spikelet fertility. BMC Plant Biol. 14, 1–13 (2014).

115. Lin, D. et al. Mutation of the rice 12 gene encoding 2,3-bisphosphoglycerate-independent phosphoglycerate mutase affects chlorophyll synthesis, photosynthesis and chloroplast development at seedling stage at low temperatures. Plant Biol. 21, 585–594 (2019).

116. Yang, L. et al. GsCBRLK, a calcium/calmodulin-binding receptor-like kinase, is a positive regulator of plant tolerance to salt and ABA stress. J. Exp. Bot. 61, 2519–2533 (2010).

117. Vega-Sánchez, M. E., Zeng, L., Chen, S., Leung, H. & Wang, G.-L. SPIN1, a K Homology Domain Protein Negatively Regulated and Ubiquitinated by the E3 Ubiquitin Ligase SPL11, Is Involved in Flowering Time Control in Rice. Plant Cell 20, 1456–1469 (2008).

118. Bentolila, S., Alfonso, A. A. & Hanson, M. R. A pentatricopeptide repeat-containing gene restores fertility to cytoplasmic male-sterile plants. Proc. Natl. Acad. Sci. U. S. A. 99, 10887–10892 (2002).

119. Nakagawa, M., Shimamoto, K. & Kyozuka, J. Overexpression of RCN1 and RCN2, rice TERMINAL FLOWER 1/CENTRORADIALIS homologs, confers delay of phase transition and altered panicle morphology in rice. Plant J. 29, 743–750 (2002).

120. He, P. et al. Identification of a fungal cytochrome P450 with steroid two-step ordered selective hydroxylation characteristics in Colletotrichum lini. J. Steroid Biochem. Mol. Biol. 220, 106096 (2022).

121. Clement, C. R. 1492 and the loss of amazonian crop genetic resources. I. The relation between domestication and human population decline. Econ. Bot. 53, 188–202 (1999).

122. Dempewolf, H., Rieseberg, L. H. & Cronk, Q. C. Crop domestication in the Compositae: A family-wide trait assessment. Genet. Resour. Crop Evol. 55, 1141–1157 (2008).

123. Hammer, K. & Khoshbakht, K. A domestication assessment of the big five plant families. Genet. Resour. Crop Evol. 62, 665–689 (2015).

